# IFN-λ4 may contribute to HCV persistence by increasing ER stress and enhancing IRF1 signaling

**DOI:** 10.1101/2020.10.28.359398

**Authors:** Olusegun O. Onabajo, Fang Wang, Mei-Hsuan Lee, Oscar Florez-Vargas, Adeola Obajemu, Mauro A. A. Castro, Chizu Tanikawa, Joselin Vargas, Shu-Fen Liao, Ci Song, Yu-Han Huang, Chen-Yang Shen, A. Rouf Banday, Thomas R. O’Brien, Zhibin Hu, Koichi Matsuda, A. Gordon Robertson, Ludmila Prokunina-Olsson

## Abstract

Chronic hepatitis C virus (HCV) infection and cirrhosis are major risk factors for developing hepatocellular carcinoma (HCC). Genetic polymorphisms in the *IFNL3/IFNL4* locus have been associated both with impaired clearance of HCV and protection from liver fibrosis, an early stage of cirrhosis. Here, we aimed to address the genetic and functional relationships between *IFNL3/IFNL4* polymorphisms, HCV-related cirrhosis, and HCC risk. We evaluated associations between *IFNL4* genotype, defined as the presence of rs368234815-dG or rs12979860-T alleles, with cirrhosis and HCC risk in patients with chronic HCV - 2,931 from Taiwan and 3,566 from Japan. We detected associations between *IFNL4* genotype and decreased risk of cirrhosis (OR=0.66, 95%CI=0.46-0.93, P=0.018, in Taiwan), but increased risk of HCC (OR=1.28, 95%CI=1.07-1.52, P=0.0058, in Japan). *In-vitro*, IFN-λ4 expression increased ER stress, and enhanced positive regulation of IFN responses via IRF1 induction, which mediated antiproliferative effects in hepatic cells. Our data present novel IFN-λ4-associated pathways that may be contributing to HCV persistence and development of HCC.

## INTRODUCTION

After being infected with hepatitis C virus (HCV), 75-85% of individuals fail to clear the virus and develop chronic infection; 10-20% of these patients progress to liver cirrhosis and to hepatocellular carcinoma (HCC), the main type of primary liver cancer, with a rate of 1-4% per year(1). Genetic polymorphisms within the *IFNL3*/*IFNL4* genomic region have been identified as the strongest predictors of spontaneous and treatment-induced clearance of HCV infection(2-5). Among these, a dinucleotide polymorphism, rs368234815-TT/dG, is the most informative marker for the prediction of HCV clearance in all populations(6, 7). Individuals with the rs368234815-dG allele have a frameshift in the first exon of the *IFNL4* gene, which creates an open reading frame for interferon lambda 4 (IFN-λ4), a type III IFN. Homozygous carriers of the rs368234815-TT allele, including most Asians (∼90%) and Europeans (∼50%), but only a minority of individuals of African ancestry (∼10%), are natural IFN-λ4 knockouts(7).

Carriers of the *IFNL4*-dG allele are more likely to develop chronic HCV infection and fail to respond to HCV treatments. Given that chronic HCV infection is a strong risk factor for HCC, the *IFNL4*-dG allele would be expected to be associated with the risk of HCC as well. A recent study of 1866 HCC patients from Biobank Japan showed a significant association between *IFNL3/IFNL4* polymorphisms and HCC risk(8), with several smaller studies making similar but not as statistically significant observations (9-13). However, the same genetic variants associated with poor HCV clearance also showed association with reduced risk of liver fibrosis(14, 15), which is an aberrant wound-healing response to chronic liver injury and a pre-stage of liver cirrhosis(16). The complexity of the associations of these polymorphisms with established HCC risk factors warrants the need for a thorough elucidation of biological mechanisms underlying these associations. Identifying these mechanisms could also reveal new pathways leading to development of HCC. Here, we evaluated how IFN-λ4 could be contributing to HCC using genetic, genomic, and functional tools.

## RESULTS

### *IFNL4* genotype is associated with increased risk of HCC in patients without HCV clearance

We analyzed the clinical REVEAL II cohort from Taiwan(17), which included 2,931 HCV-infected patients who were treated with peg-IFNα/RBV and then monitored for progression to HCC (**Table S1**). Due to strong linkage disequilibrium (LD) in the *IFNL3*/*IFNL4* region(7), many markers provide similar genetic associations, including rs368234815 that controls IFN-λ4 production(6), rs4803217 that was suggested to affect IFN-λ3 stability(18), and rs12979860, a non-functional polymorphism in intron 1 of *IFNL4*(3). We genotyped *IFNL4*-rs368234815, which has a direct functional effect on IFN-λ4 production(6, 19). Since the rs368234815-dG allele (which supports the production of IFN-λ4) is uncommon in Asian populations, we used a dominant genetic model and combined all carriers of the dG allele (TT/dG and dG/dG) in one group, designated as the *IFNL4* genotype group.

As expected(2, 4), carriers of *IFNL4* genotype had reduced sustained virologic response (SVR) after treatment or retreatment with IFNα/RBV (p=1.63E-18, **Table S2**), but we also observed a lower prevalence of cirrhosis at baseline (before treatment, p=0.018, **Table 1**). Among all REVEAL II patients, *IFNL4* genotype was associated with progression to HCC (adjusting for cirrhosis, p=0.033, **Table S3**), but this association was no longer significant after accounting for SVR after treatment or retreatment (**Table S3, S4**).

**Table 1.** Prevalence of cirrhosis in HCV-infected patients in relation to *IFNL4* genotype in the REVEAL II cohort, Taiwan

| Genotype of<br>*IFNL4*-<br>rs368234815 | Liver cirrhosis at baseline,<br>N=2931<br>N (%) |  | OR (95% CI), P-value |  |
| --- | --- | --- | --- | --- |
|  | Yes<br>N=508<br>N (%) | No<br>N=2403<br>N (%) | Crude | adjusted for<br>age, sex |
| TT/TT | 468 (17.88) | 2150 (82.12) | Ref | Ref |
| TT/dG or dG/dG | 40 (12.78) | 273 (87.22) | 0.67<br>(0.48-0.95)<br>P=0.025 | 0.66<br>(0.46-0.93)<br>P=0.018 |

TT/TT genotype – IFN-λ4 is not produced

We also analyzed HCC risk in 3,566 HCV-infected patients from the BioBank Japan(20, 21), using an intronic *IFNL4* marker rs12979860, which is completely linked with rs368234815 in Asians (r^2^=1.0). The *IFNL4*-rs12979860-T allele was associated with an increased risk of HCC (p=0.0058), both in patients with (p=0.047) and without cirrhosis (p=0.011, **Table 2**). Because we selected patients based on their HCV-positive status, this association represents patients who have failed to clear HCV either spontaneously or after treatment, but these questions were not explored due to lack of information on treatments and outcomes. *IFNL4* genotype was not associated with progression to HCC in Chinese patients with HBV (**Table S5**). These results demonstrate the association of *IFNL4* genotype with increased risk of HCC, despite a modest association with a lower incidence of liver cirrhosis. These results also emphasize the critical importance of achieving HCV clearance for preventing the progression to HCC.

**Table 2.**
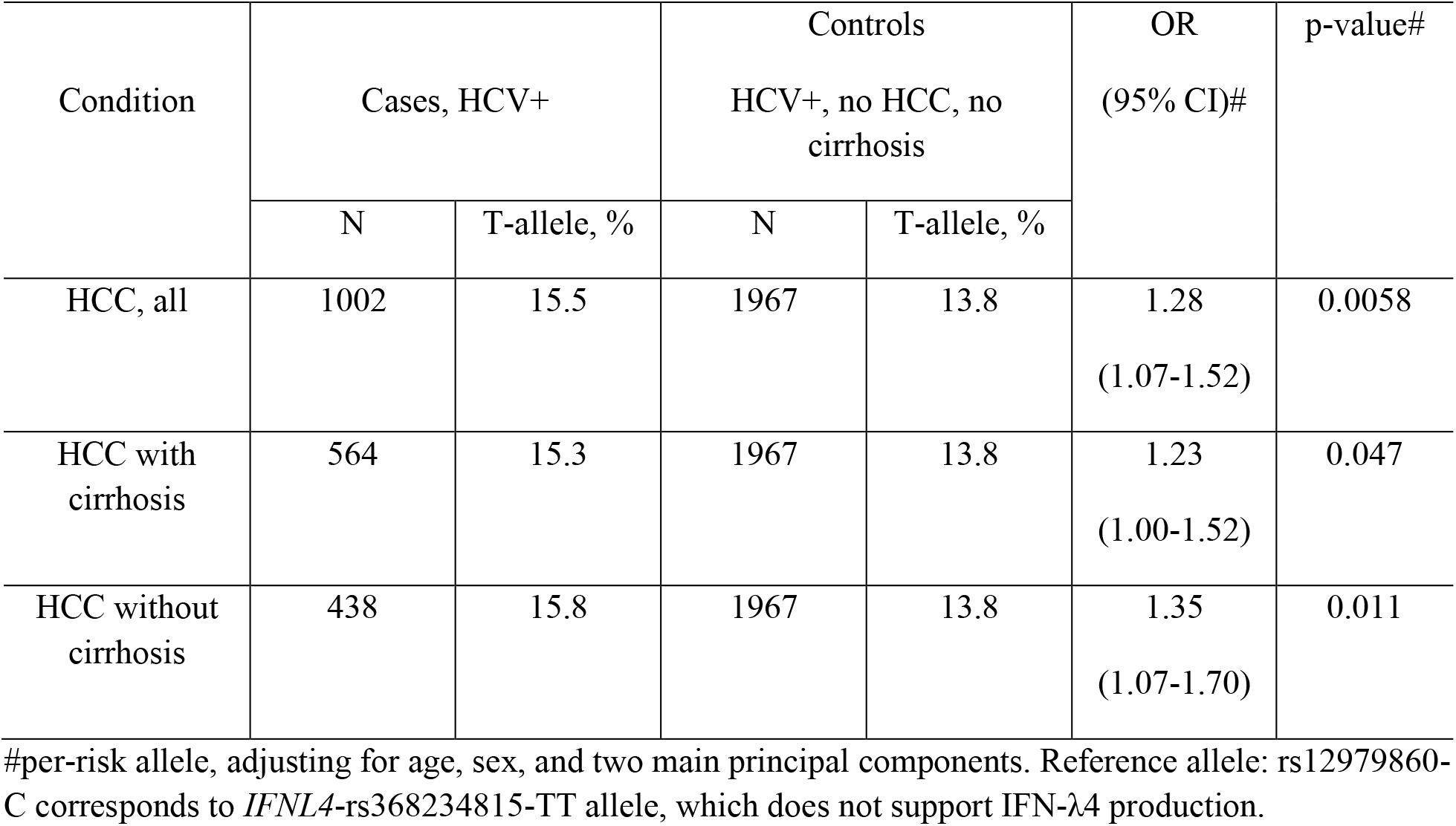
*IFNL4* genotype is associated with increased risk of HCC in HCV-infected patients in Japan

| Condition | Cases, HCV+ |  | Controls<br>HCV+, no HCC, no<br>cirrhosis |  | OR<br>(95% CI)# | p-value# |
| --- | --- | --- | --- | --- | --- | --- |
|  | N | T-allele, % | N | T-allele, % |  |  |
| HCC, all | 1002 | 15.5 | 1967 | 13.8 | 1.28<br>(1.07-1.52) | 0.0058 |
| HCC with<br>cirrhosis | 564 | 15.3 | 1967 | 13.8 | 1.23<br>(1.00-1.52) | 0.047 |
| HCC without<br>cirrhosis | 438 | 15.8 | 1967 | 13.8 | 1.35<br>(1.07-1.70) | 0.011 |

#per-risk allele, adjusting for age, sex, and two main principal components. Reference allele: rs12979860-C corresponds to *IFNL4*-rs368234815-TT allele, which does not support IFN-λ4 production.

### IFN-λ4 expression induces cell cycle arrest and inhibits proliferation of hepatic cells

Next, we aimed to explore molecular mechanisms underlying the association between *IFNL4* genotype and increased risk of HCV persistence. Expression and function of both IFN-λ3 and IFN-λ4 might be affected by the associated genetic variants, making it hard to delineate the individual contribution of these IFNs by genetic analysis alone. Thus, we explored functional effects of both IFN-λ3 and IFN-λ4 in a set of hepatoma HepG2 cell lines, in which we inducibly expressed IFN-λ3-GFP or IFN-λ4-GFP, and used CRISPR-Cas9 gene editing to eliminate IFNLR1, the receptor used by all type III IFNs (**Fig. S1, Fig. S2**). After doxycycline (dox)-induction of these cell lines for 8, 24 and 72 hrs, we analyzed their global transcriptome by RNA-seq. IFN-λ4 induction for 72 hrs resulted in the largest set of differentially expressed genes (DEGs, P-FDR < 0.05 and dox+/dox-fold change +/-≥ 1.5, n=2880); this set was used for further comparative analyses (**Fig. S3, Table S6**). Of these genes, 2,735 and 145 were classified as IFNLR1-dependent and-independent, respectively, based on their differential expression in IFN-λ4-GFP and IFN-λ4-GFP-IFNLR1^KO^ cells (**Fig. 1A**). The set of IFN-λ4-induced and IFNLR1-dependent genes (n=2,735) was then compared with the expression of the same gene set in the IFN-λ3 transcriptome. *IFNL3* and *IFNL4* transcripts in these cell lines were induced to a similar magnitude, making these results comparable (**Fig. S4**). A total of 1,506 genes were induced by both IFN-λ3 and IFN-λ4 (P-FDR < 0.05), with 766 genes induced by ≥1.5 fold in both groups. As expected, ISGs were the top-induced genes, which were activated faster and stronger by IFN-λ4 compared to IFN-λ3 (**Fig. S5, Fig. 1B**). The remaining 1,229 (of 2,735) genes, which we considered as IFN-λ4-specific DEGs (**Fig. S3G, Table S5**), were enriched with genes related to inhibition of cell cycle (**Fig. S6**), consistent with IPA results in TCGA-LIHC (**Fig. 1C**).

**Fig. 1.**
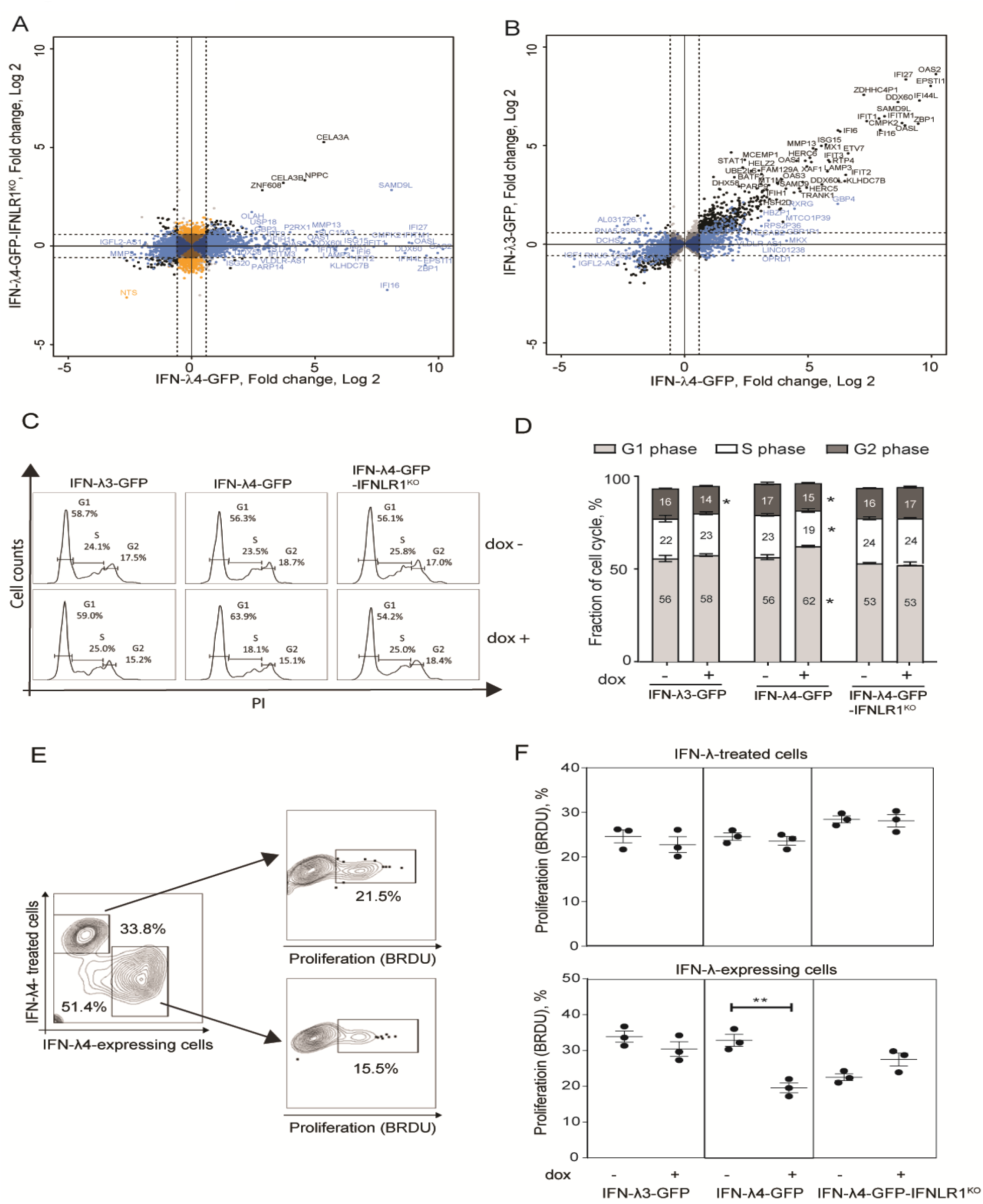
IFN-λ4 overexpression inhibits the proliferation of hepatic cells by inducing cell cycle arrest. Differentially expressed genes (DEGs, P-FDR < 0.05) were identified by RNA-seq analysis in IFN-λ3-GFP, IFN-λ4-GFP and IFN-λ4-GFP-IFNLR1^KO^ HepG2 cells after 72 hrs of induction by dox, comparing to controls (dox-conditions). Cutoff threshold (fold change > +/-1.5) is indicated by dotted lines. **(A)** Analysis of all DEGs (n=3251) detected for IFN-λ4-GFP or IFN-λ4-GFP-IFNLR1^KO^ cells. In blue - DEGs (n=2,735) specific to IFN-λ4-GFP and considered IFNLR1-dependent. In black - DEGs (n=145) shared between both groups and considered IFNLR1-independent. In orange - DEGs (n=371) specific to IFN-λ4-GFP-IFNLR1^KO^. **(B)** DEGs of IFN-λ4-GFP analyzed in IFN-λ3-GFP transcriptome. In black - DEGs (n=1,506) shared in IFN-λ4-GFP and IFN-λ3-GFP and in blue - IFN-λ4-signature DEGs (n=1,229) detected in IFN-λ4-GFP but not in IFN-λ3-GFP producing cells. Additional details are provided in **Fig. S3** and **Table S6. (C**,**D)** Cell cycle analysis of cells synchronized by 24 hrs of serum starvation, treated with or without dox (0.5 µg/ml) for 72 hrs and analyzed by flow cytometry after PI staining. The plot shows a representative picture and the percentage of cells in each phase of the cell cycle. All data are shown as mean± SEM from triplicate experiments. *, P < 0.05. **(E**,**F)** Bromodeoxyuridine (BRDU, %) incorporation indicating cell proliferation in HepG2 cells expressing IFN-λ3-GFP, IFN-λ4-GFP and IFN-λ4-GFP-IFNLR1KO. Cells were cocultured with HepG2 cells labeled with Far Red proliferation dye, dox-induced for 72 hrs and treated with BRDU for 3 hrs before analysis. Gates show HepG2 cells exposed to IFN-λs (IFN-λ treated cells) and HepG2 expressing IFN-λs. P-values compare corresponding dox+ vs. dox-HepG2 cells, ** p<0.01, Student’s T-test. Graphs represent one of three independent experiments, each in biological triplicates.

In line with our previous results(22), in IFN-λ4-expressing HepG2 cells, we observed G1/G0 cell cycle arrest (**Fig. 1C, D**) and reduced proliferation (**Fig. S7**). However, the antiproliferative effect was absent when HepG2 cells were cocultured with IFN-λ4-expressing cells or when IFN-λ4 was overexpressed in IFNLR1^KO^ cells (**Fig. 1E,F**). These results indicate that IFN-λ4 causes cell cycle arrest and decreased proliferation of hepatic cells through a mechanism that is IFNLR1-dependent but requires intracellular expression of IFN-λ4.

### IFN-λ4 inhibits proliferation of hepatic cells by inducing IRF1

We then assessed our curated list of DEGs generated in HepG2 cells (**Fig. S3**) in relation to regulatory networks of 885 transcription factors and their targets (“regulons”(23, 24)), which we constructed for 373 TCGA-LIHC tumors (see methods). Of the 885 TCGA-LIHC regulons with at least ≥15 targets, 54 regulons were significantly and directionally enriched with HepG2-DEGs (**Table S7**). Unsupervised clustering of regulon activity profiles using HepG2-DEGs identified groups of regulons with low vs. high activity scores (**Fig. 2A**). Group I included regulons activated by IFN-λ4, such as STAT1 and IRF9 (**Fig. 2A**), both of which are critical for the receptor-dependent signaling of type III IFNs(25), thus validating our analysis. Group II included regulons that were repressed by IFN-λ4, such as CENPA, FOXM1, and TET1 (**Fig. 2A**), that are transcription factors associated with cell cycle progression, consistent with our IPA results.

**Fig. 2.**
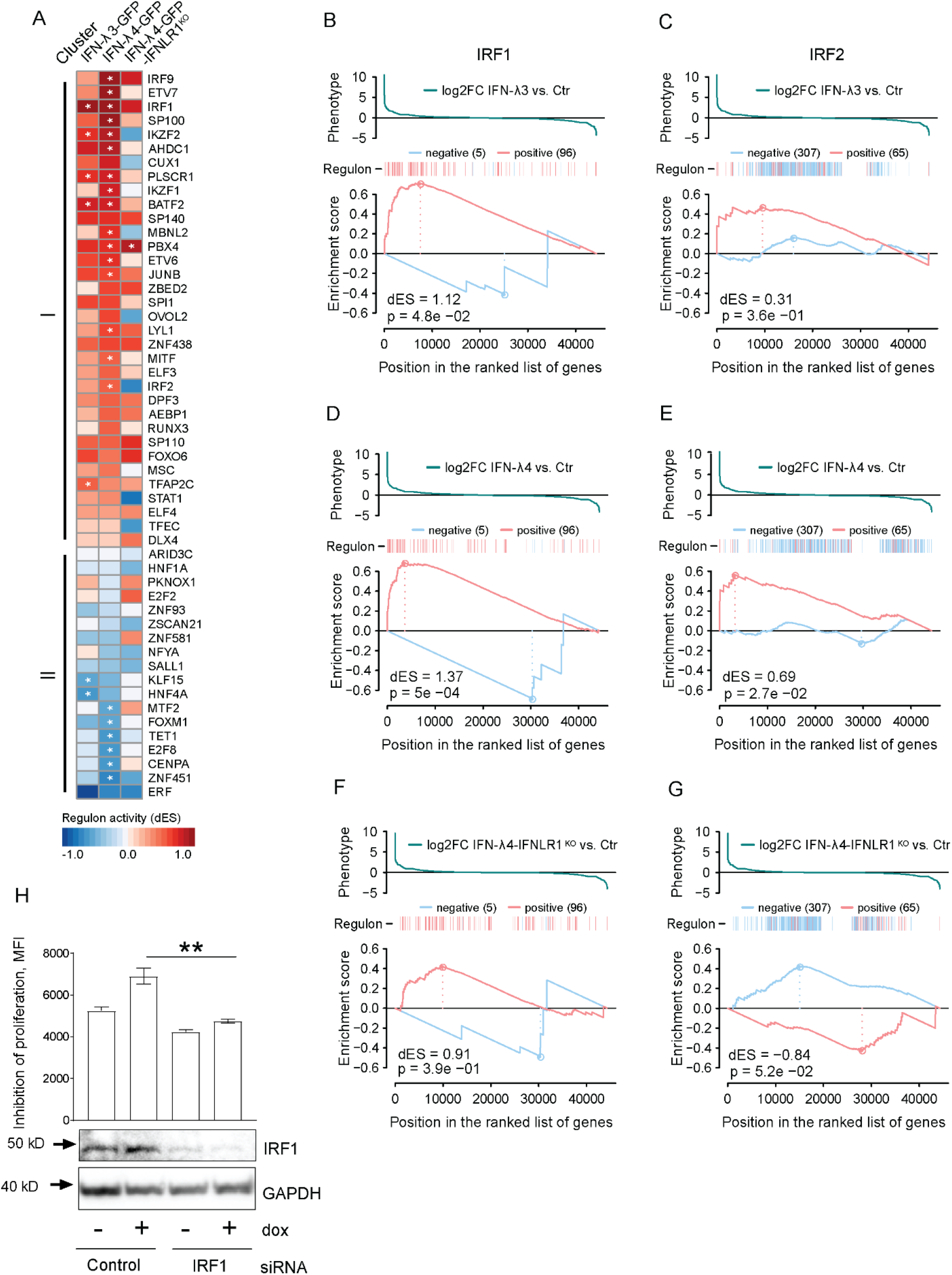
IRF1 is a critical IFN-λ4-induced regulator of cell proliferation. **(A)** Unsupervised clustering of activities of 54 regulons that were significantly and directionally enriched with HepG2-DEGs in the set of 885 of TCGA-LIHC regulons. The heatmap shows differences in activity scores (dES) for the IFN-λ4-enriched regulons organized by GSEA-2T results for the IFN-λ4-GFP, IFN-λ3-GFP and IFN-λ4-GFP-IFNLR1^KO^ DEG signatures. Cluster I: regulons with dES > 0 in IFN-λ4-GFP; cluster II: regulons with dES < 0 in IFN-λ4-GFP. **Table S7** provides the regulon activity scores presented in **Figure 2A. (B-G)** GSEA-2T plots for IRF1 and IRF2, respectively, in DEG signatures for **(B-C)** IFN-λ3-GFP, (**D-E**) IFN-λ4-GFP, and (**F-G**) IFN-λ4-GFP-IFNLR1^KO^. **(H)** Inhibition of proliferation in IFN-λ4-GFP HepG2 cells, for one of three independent experiments. Cells were treated with IRF1 siRNA for 24 hrs, labeled with Far Red proliferation dye and dox-induced at indicated concentrations for 72 hrs. Proliferation was assessed by flow cytometry with a graph representing the geometric mean expression of Far Red proliferation dye with higher values indicating reduced cell proliferation. P-values compare dox-treated control siRNA with dox-treated IRF1 siRNA. ** p<0.01, Student’s T-test. Below: Western blots showing IRF1 protein levels following siRNA knockdown.

The regulon of interferon regulatory factor 1 (IRF1) was strongly activated by IFN-λ4 expression (**Table S7**). IRF1 is a transcription factor that enhances the antiviral response but reduces liver cirrhosis by inhibiting proliferation of hepatic stellate cells (HSCs)(26, 27); IRF1 activity is antagonized by the closely related IRF2(28). Gene set enrichment analysis (GSEA)(29) showed that the IRF1 regulon enrichment scores were higher for IFN-λ4, compared to IFN-λ3 and IFNLR1^KO^ (**Fig. 2B-G**), and *IRF1* expression was significantly induced by IFN-λ4 and to a lesser extent by IFN-λ3 (**Fig. S8**). The IFN-λ4-induced activity of IRF2 regulon was weak (**Fig. 2E)** and unlikely to impact IRF1 regulon activity. Importantly, the antiproliferative effect of IFN-λ4 in HepG2 cells was attenuated by *IRF1* knockdown (**Fig. 2H**), supporting the critical role of IRF1 in this mechanism.

### IFN-λ4 is a misfolded protein that induces a potent ER stress response in hepatic cells

IFN-λ4-producing cells had no antiproliferative effect on bystander cells (**Fig. 1E, F**), suggesting that paracrine IFN signaling alone was insufficient to inhibit cell proliferation, prompting us to search for additional intrinsic factors associated with IFN-λ4 expression. Unfolded protein response (UPR) was one of the main pathways identified by IPA as significantly activated in IFN-λ4-expressing HepG2 cells (**Fig. S9**). UPR due to endoplasmic reticulum (ER) stress has been shown to inhibit proliferation and induce apoptosis in some conditions(30, 31). UPR is induced to relieve ER stress by increasing the expression of several chaperones that aid protein folding(30, 32), and we found a stronger induction of UPR effectors in IFN-λ4-expressing compared to IFN-λ3-expressing and IFN-λ4-IFNLR1^KO^ HepG2 cells (**Fig. S10**). We previously showed that infection of primary human hepatocytes (PHH) with Sendai virus (SeV) resulted in intracellular accumulation of IFN-λ4(19), possibly causing ER stress. Now, we tested this hypothesis and confirmed the upregulation of ER stress genes in PHH from donors with vs. without *IFNL4* genotype (**Fig. S11**).

The induction of ER stress and poor secretion of IFN-λ4 due to its intracellular accumulation(19, 22) suggested that IFN-λ4 is an inefficiently folded protein that induces the misfolded protein ER stress response(31, 33). To mitigate ER stress, misfolded proteins are targeted to lysosomes for degradation(30, 33). Live imaging in HepG2 cells showed that IFN-λ4 was accumulated in lysosomes (**Fig. 3A**) via late endosome trafficking (**Fig. 3B, Video S1**) but was excluded from early recycling endosomes (**Fig. 3A**). Continued accumulation of IFN-λ4 led to lysosomal enlargement (**Fig. 3C**), followed by membrane blebbing and cell death, implicating apoptosis (**Fig. 3D, Video S2**). We confirmed apoptosis in IFN-λ4-expressing cells using a biochemical assay for caspase activity (**Fig. 3E**) and cell viability (**Fig. 3F**). Prolonged ER stress causes apoptosis via the activity of the UPR-induced transcription factor DDIT3(34), which downregulates the expression of antiapoptotic factors, including BCL2(35) (**Fig. S9**). This led us to postulate that blocking the *DDIT3* expression might attenuate the pro-apoptotic effects of IFN-λ4. Indeed, reduction of *DDIT3* expression by RNA interference in IFN-λ4-expressing cells (**Fig. 3G, H**) resulted in inhibition of cell apoptosis by 20% (**Fig. 3I**) and improved cell viability by 30% (**Fig. 3J**). These results indicate that prolonged ER stress due to the continued endogenous production of misfolded IFN-λ4 could be contributing to apoptosis of hepatic cells.

**Fig. 3.**
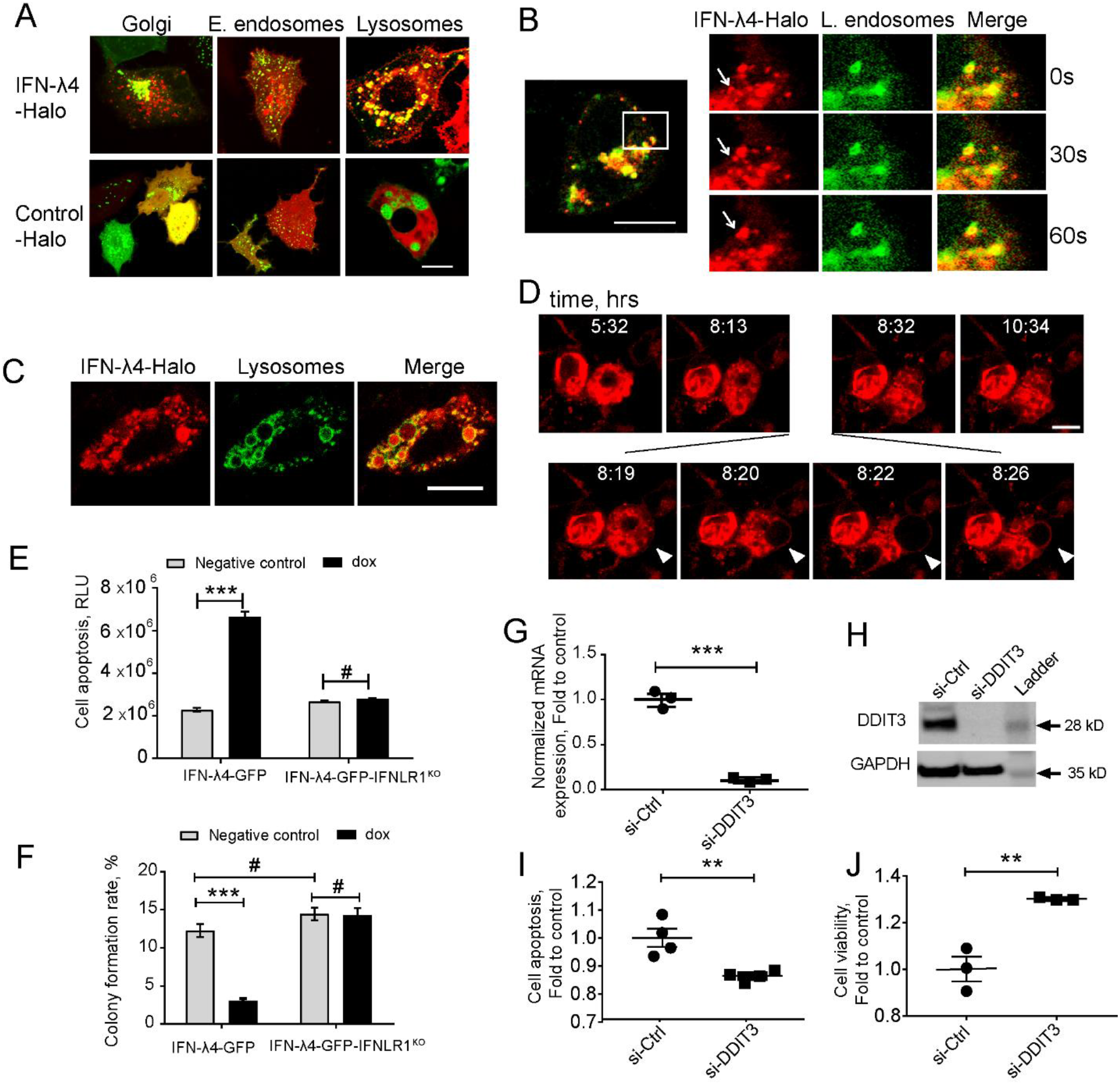
IFN-λ4 is a misfolded protein that induces ER stress. Representative confocal images of HepG2 cells transduced with a mammalian baculovirus delivery system (BacMam) of GFP-tagged proteins targeting specific organelles - lysosomes, Golgi, early and late endosomes. After transduction for 6 hrs, cells were transiently transfected with Halo-tagged constructs for IFN-λ4 or control for indicated times, stained with cell-permeant Halo-tag ligand TMR (red), and imaged. **(A)** Confocal images showing IFN-λ4 accumulation in lysosomes but not in early endosomes. **(B)** Late endosomal trafficking of IFN-λ4, with the inset showing larger magnification. **(C)** Unfolded protein response (UPR) is represented by lysosomal enlargement after protein accumulation. **(D)** Live images of IFN-λ4-expressing HepG2 cells undergoing apoptosis, characterized by membrane blebbing and cell death. Images were scanned every minute for 12 hrs. Scale bars – 10 um. **(E)** Apoptosis detection with ApoTox-Glo assays in corresponding untreated and dox-induced cells for indicated time points. RLU, relative luminescence units. **(F)** Graph showing counts from colony formation assay for HepG2 cells expressing IFN-λ4 or IFNLR1 KO grown in 6-well plates with or without dox for 13 days. Cell colonies were stained with crystal violet and manually counted. The graph represents the number of colonies as a percentage of initial plated counts. **(G,H)** mRNA **(G)** and protein levels (**H**) of DDIT3 after siRNA knockdown tested by qRT-PCR and Western blot assays, respectively. **(I-J)** Apoptosis **(I)** and cell viability **(J)** assays were performed after siRNA knockdown of DDIT3 in dox-induced IFN-λ4-GFP cells. * p<0.05, ** p< 0.01, *** p<0.001.

### IFN-λ4 induces ER stress in hepatic stellate cells

Activated hepatic stellate cells (HSCs) are key drivers of liver fibrosis during HCV infection(36), and while HCV is not known to replicate in HSCs, HSCs can produce interferons in response to HCV(37). Thus, we examined whether IFN-λ4 can induce ER stress not only in hepatocytes but also in HSCs. We infected primary HSCs with SeV to induce IFNs and observed significant induction both of *IFNL4* (**Fig. 4A**) and ER stress genes (**Fig. 4B**). The induction of *OAS1*, an ISG, in response to treatment with IFN-λ4 indicated the presence of the functional type III signaling machinery, including IFNLR1, in primary HSCs (**Fig. 4C**). When we overexpressed IFN-λ4 in LX2 (an HSC cell line), similarly to hepatocytes, we observed intracellular accumulation of IFN-λ4, causing upregulation of ER stress genes, and inhibition of cell proliferation (**Fig. 4D-G**). These results suggest that IFN-λ4 expression during HCV could mitigate the aberrant proliferation of HSCs and slow down the development of fibrosis.

**Fig. 4.**
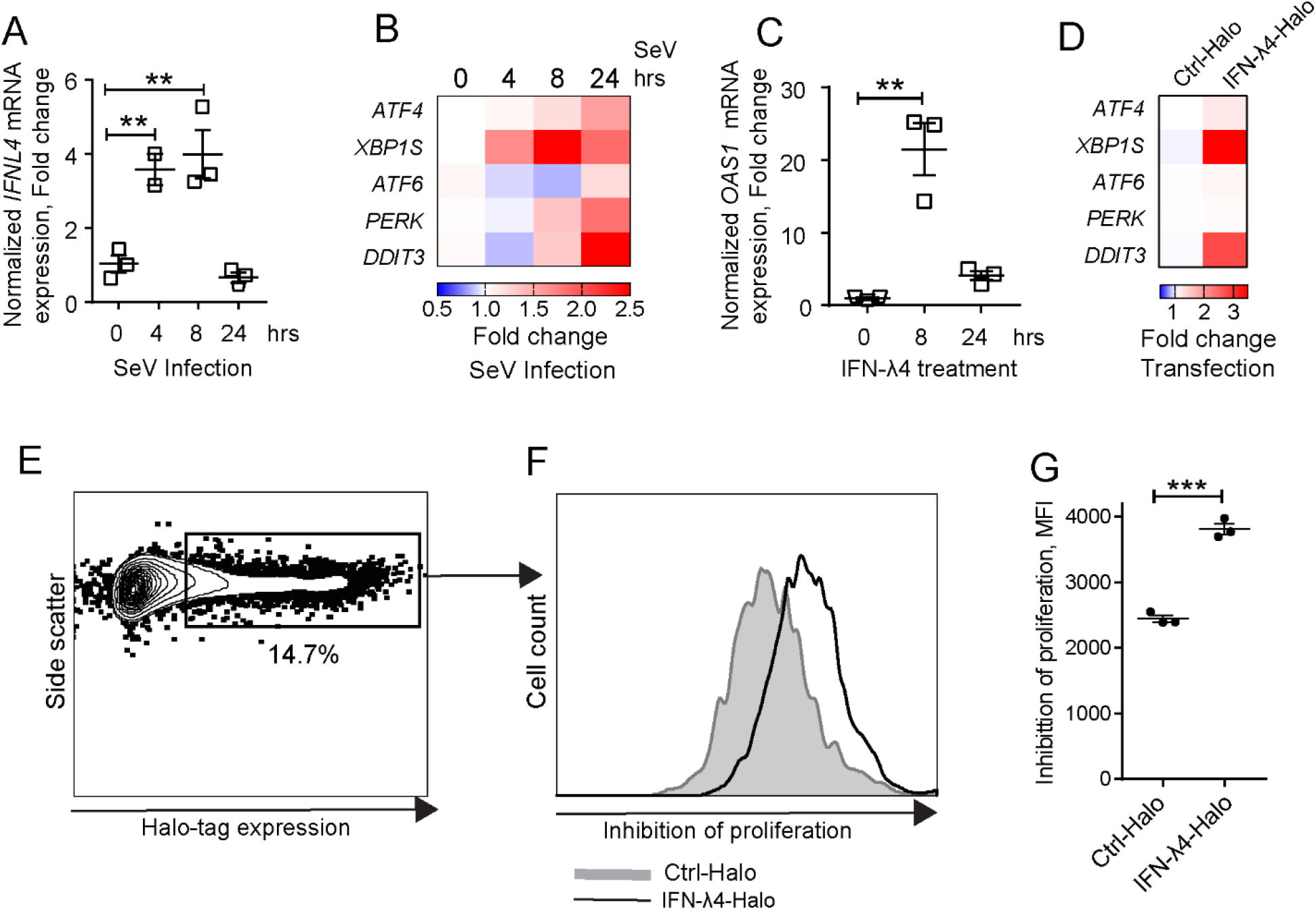
IFN-λ4 inhibits the proliferation of hepatic stellate cells (HSCs). qRT-PCR expression analysis showing **(A)** *IFNL4* expression after SeV infection of primary HSCs and **(B)** induction of UPR markers. **(C)** qRT-PCR showing induction of *OAS1* after treatment of primary HSCs with recombinant IFN-λ4 (50ng/ml). **(D)** qRT-PCR showing induction of UPR markers in LX-2 hepatic stellate cell line transiently transfected for 24 hrs with Halo-tagged constructs for IFN-λ4 or control. (**E-G**) Proliferation analysis in LX-2 cells transiently transfected with Halo-tagged constructs for IFN-λ4 or control, labeled with Far Red proliferation dye and analyzed by flow cytometry after 72 hrs. (**E)** Gating for Halo-tag expression. **(F)** Histogram representing the proliferation of gated cells, comparing cells expressing Halo-tagged IFN-λ4 vs. control. **(G)** Graph representing the geometric mean expression of Far Red proliferation dye with higher values indicating reduced proliferation.** p< 0.01, *** p<0.001, Student’s T-test.

## DISCUSSION

Based on our genetic analyses in large sets of patients and functional *in-vitro* studies, we conclude that IFN-λ4 plays multiple roles in cirrhosis and HCV risk. The first and most known role of IFN-λ4 is in interfering with viral clearance and sustaining chronic HCV infection, a risk factor for HCC. In our study, only patients without HCV clearance had a significant *IFNL4* genotype-associated risk of HCC. *IFNL4* genotype was also associated with reduced cirrhosis, but this effect was not sufficient to mitigate the increased risk of developing HCC. Protection from cirrhosis could be due to antiproliferative and apoptotic effects of IFN-λ4 expression in hepatic cells, particularly in stellate cells. This role required ER stress and enhanced IRF1 signaling, resulting from the intracellular accumulation of misfolded IFN-λ4 protein. It is important to emphasize that in the context of HCV, the modest anticirrhotic effects of IFN-λ4 do not appear sufficient to mitigate the association with HCC development. It is possible that the anticirrhotic effect of IFN-λ4 could be relevant for other forms of non-HCV-related hepatitis.

Viral hepatitis is characterized by repeated cycles of hepatic cell death and regeneration. Breakdown in the formation of the extracellular matrix in combination with continuous proliferative pressure leads to the development of fibrotic tissue(16, 38-40). This mechanism is mediated by HSC, which differentiate into myofibroblasts and accumulate in fibrotic tissue(40, 41). As a result, signaling pathways that lead to cell cycle arrest and apoptosis of HSC inhibit the development of fibrosis and cirrhosis(26, 40). Indeed, we observed that IFN-λ4-induced ER stress in HSC coupled with IFN signaling reduced proliferation and increased apoptosis, which could be anticirrhotic.

IFNs, including IFN-λ3 and IFN-λ4, are expected to affect the expression and activity of many transcription factors, and the networks of downstream targets they regulate. Of the 885 regulons identified in TCGA-LIHC and probed with expression signatures experimentally derived in HepG2 cells, we noted several expected and a number of novel regulons. Notably, we observed a rather weak induction of IRF1 regulon by IFN-λ3 (comparable to IFNLR1^KO^ cells), consistent with the previously reported inferior ability of type III IFNs to induce IRF1 and its proinflammatory program when compared to type I IFNs(27, 42). IFN-λ4, on the other hand, induced significantly higher levels of *IRF1* expression than IFN-λ3, resulting in stronger activation of the IRF1 regulon. IRF1 signaling was reported as an important anticirrhotic factor(26) and the antiproliferative effect of IFN-λ4 appeared to be dependent on the ability to induce IRF1 signaling. Previous studies suggested that the induction of *IRF1* expression was the main difference between the transcriptomes of cells treated with type I vs. type III IFNs(27, 42). Specifically, *IRF1* was induced strongly by treatment with IFNβ, while only weakly or not induced by treatment with IFN-λs(27, 42). Our previous(19, 22) and current results suggest that *IRF1* induction by treatment with exogenous IFN-λ4 is weaker compared to that induced by the intracellular IFN-λ4 expression, and might be insufficient for inducing relevant functional effects, including inhibiting cell proliferation.

The higher frequency of *IFNL4* genotype in African-Americans (∼70% *IFNL4*-dG allele frequency) compared to European-Americans (∼30%), and the protective association of this genotype with liver cirrhosis might explain some of the disparity in cirrhosis-related death rates reported in the US(43). During 2000-2013, these rates trended in opposite directions, increasing in European-Americans while decreasing in African-Americans(43). The rates of HCV infection and progression to HCC in these individuals are unknown, but it would be interesting to further explore the potential association of *IFNL4* genotype with these outcomes and corresponding mortality.

The reasons for the strong association of the *IFNL4* genotype with an increased risk of developing chronic HCV remain a puzzle. A recent report suggested that IFN-λ4 may be reducing our ability to fight infections broadly (44). Recipients of hematopoietic stem cell transplantation (HSC) for acute leukemia receiving the transplant from donors with *IFNL4* genotype were significantly more likely to die from post-transplant complications, including infections, compared to *IFNL4*-null donors. Surprisingly, *IFNL4* was the most frequently expressed IFN in bone marrow cells, suggesting that IFN-λ4 may be playing a more broad role in reprogramming immune response manifesting in various disease outcomes.

In conclusion, we present genetic and functional results reconciling the role of IFN-λ4 in protecting from liver cirrhosis but increasing the risk of HCC by sustaining the persistence of HCV infection. We acknowledge the limitations of our study, including the lack of animal models, complicated by the absence of *IFNL4* in the mouse genome. Dissecting the role of IFN-λ4 in pathogenic processes driving HCC in the context of HCV infection and developing clinically relevant applications will require replicating these findings in comparable cohorts and controlling for HCV treatments and outcomes.

## MATERIALS AND METHODS

### HCV+ patients from REVEAL II cohort, Taiwan

The REVEAL-II (Risk Evaluation of Viral Load Elevation and Associated Liver Diseases) clinical cohort has been described (17), (**Table S1**). Briefly, treatment-naive HCV+ patients without HCC were enrolled during 2004–2014 in the prospective study. The patients were treated with Peg-IFN and ribavirin (RBV) for 48 or 24 weeks (for patients with HCV genotype 1 and other genotypes, respectively) and sustained virologic response (SVR) was determined from the serum HCV RNA test results 24 weeks post-treatment. Patients who didn’t achieve an SVR were re-treated with the same regimen and the combined SVR was estimated based on the treatment and retreatment.

Demographic and clinical data were obtained via chart reviews with standardized forms. Cirrhosis at baseline (before treatment) was determined by abdominal ultrasound (in 81.6% of patients) or liver biopsy (in 18.4%), and categorized as yes/no. DNA samples were genotyped for *IFNL4*-rs368234815 by a custom TaqMan genotyping assay, as previously described (6). Only individuals with available information about baseline cirrhosis and *IFNL4*-rs368234815 genotypes were included in the analysis (n=2,931). Incident HCC diagnoses after treatment completion were verified by medical records and electronic linkage with the Taiwan National Cancer Registration and the National Death Certification databases.

The person-years of follow-up for each patient were calculated from the enrolment date to either the date of HCC identification, the date of death, or December 31, 2014, whichever came first. The incidence rates of HCC per 1,000 person-years were calculated by dividing the number of newly developed HCC cases by person-years of follow-up. Multivariable Cox proportional hazards models were used to examine the outcomes in relation to *IFNL4* genotypes (as 0 and 1), controlling for several clinical predictors including age, sex, baseline liver cirrhosis, serum alanine aminotransferase (ALT), HCV genotype (genotype 1 vs. other), and SVR. Hazard ratios (HRs) with 95% confidence intervals (CIs) and statistical significance levels were determined by a two-sided p-value of 0.05. The proportionality assumption of Cox models was examined, and the assumption was not violated. All analyses were performed using the SAS statistical software package (version 9.1; SAS Institute Inc., Cary, NC, USA).

### HCV+ patients from BioBank Japan

A total of 3,566 individuals were from BioBank Japan, which between 2003 and 2007 recruited ∼200,000 patients with 47 common diseases from 12 Japanese medical institutes representing 67 hospitals (20). From the BioBank Japan database, we selected all the patients who were HCV-RNA positive at the time of recruitment, had cirrhosis status determined by ultrasound (yes/no) and were ascertained for primary HCC status (yes/no) based on histology, imaging, and laboratory tests. No information about HCV treatment and outcomes was available. The project was approved by the ethical committees of the University of Tokyo and all participants provided written informed consent. Genomic DNA from peripheral blood was genotyped with an Illumina HumanOmniExpressExome BeadChip or a combination of the Illumina HumanOmniExpress and HumanExome BeadChips (21). The genotypes were prephased with MACH (45) and imputed with Minimac using the 1000 Genomes Project Phase 1 (version 3) East Asian reference panel (46). *IFNL4*-rs12979860 was imputed with high confidence and association analysis was conducted using a logistic regression model using imputed gene dosage, adjusting for age, sex, and the top 2 principal components (PCs).

### HBV+ patients from China

HBV-positive HCC patients (n=1,300) were consecutively recruited between 2006 and 2010 in Nantong and Nanjing, China (47). Age and sex-matched controls were recruited from the same geographical areas and included 1,344 HBV persistent carriers that were positive for both HBsAg and antibody against hepatitis B core antigen (anti-HBc), as well as 1,344 subjects with natural clearance of HBV - negative for HBsAg, but positive for antibodies against hepatitis B surface antigen (anti-HBs) and anti-HBc. All the cases and controls were negative for HCV antibody (anti-HCV). No information about HBV treatment and outcomes was available.

Genomic DNA was extracted from blood leukocytes with phenol-chloroform. DNA samples were genotyped for *IFNL4*-rs368234815 by a custom TaqMan genotyping assay as previously described (6). Association analysis was conducted using a logistic regression model adjusting for relevant covariates.

### Human cells lines and primary cells

Human cell lines: HepG2 (hepatoma) and 293T (embryonal kidney) were acquired from the American Type Culture Collection (ATCC), while LX-2 (hepatic stellate) was purchased from Millipore-Sigma. Stable HepG2 cell lines expressing doxycycline (dox)-inducible GFP-tagged IFN-λ3 and IFN-λ4 (IFN-λ3-GFP and IFN-λ4-GFP cells, respectively, have been described (19). All cell lines were maintained in DMEM supplemented with 10% FBS (Invitrogen), penicillin, and streptomycin; 5 μg/ml blasticidin and 1 mg/ml neomycin were additionally used for the stable cell lines. Expression of IFN-λ3–GFP and IFN-λ4–GFP was induced by 0.5 μg/ml or specified concentrations of dox for indicated time points. LX-2 cells were maintained in 2% FBS, 100 units/ml penicillin, 100 µg/ml streptomycin and 2 nM glutamine media. Primary human hepatic stellate cells (HSCs) were purchased from ScienCell Research Laboratories and maintained in stellate cell media with 10% FBS and stellate cell growth supplement (ScienCell). All cell lines were tested bi-monthly for mycoplasma contamination using the MycoAlert Mycoplasma Detection kit (Lonza) and were authenticated annually using the AmpFLSTR Identifiler Plus Kit (ThermoFisher Scientific) by the Cancer Genomics Research Laboratory (CGR, NCI). Fresh primary human hepatocytes (PHHs) from 12 anonymous donors purchased from BioreclamationIVT and genotyped for *IFNL4*-rs368234815 were previously described (19).

### Generation of IFNLR1 knockout cell line using CRISPR/Cas9 genome editing

Exon 3 of the NM_170173.3 transcript, which is the first exon common for all four alternative isoforms of *IFNLR1*, was selected as a target region for designing six candidate gRNAs with a CRISPR design tool (http://crispr.mit.edu/). Double-stranded oligonucleotides were cloned into the linearized SmartNuclease vector (EF1-T7-hspCas9-T2A-RFP-H1-gRNA with red fluorescent protein (RFP, System Biosciences) and IFNLR1-guide RNA (IFNLR1-gRNA) plasmids were validated by DNA sequencing. 293T cells were transiently transfected in 6-well plates with 6 individual IFNLR1-gRNA plasmids using Lipofectamine 3000 (Life Technologies). The cells were harvested 4 days post-transfection and subjected to fluorescence-activated cell sorting (FACS) for RFP with a FACS Aria III (BD Biosciences). Genomic DNA from sorted RFP-positive cells was extracted using DNeasy Blood & Tissue Kit (Qiagen) and used for sequencing of the exon 3 target region. One out of 6 IFNLR1-gRNA plasmids was selected as the most optimal (**Fig. S1A**) and was transfected into IFN-λ4-GFP cells in a 12-well plate, together with linearized screening puromycin marker (Clontech). Two days post-transfection, cells were plated in a 10-cm plate and fresh media containing puromycin (2 µg/ml) was added to the plate two days later. After three weeks of selection, visible clones were picked and transferred into 24-well plates with fresh media without puromycin. DNA was extracted from each of the clones and sequenced as described above. Final clones represented the stable IFN-λ4-GFP-IFNLR1^KO^ cells. Potential off-target sites were predicted with the online tool CCTop (http://crispr.cos.uni-heidelberg.de) (48). Top ten predicted sites were tested by sequencing of the genomic DNA of IFN-λ4-GFP-IFNLR1^KO^ cells and no off-target mutations were detected (data not shown).

### Interferon-stimulated response element luciferase reporter (ISRE-Luc) assays

Cells were seeded in 96-well plates (2 × 10^4^ cells/well) and reverse-transfected with 100 ng/well of Cignal ISRE-Luc reporter (Qiagen) using Lipofectamine 3000. After 24 hours of transfection, cells were treated for 8 hours in triplicates with recombinant IFNα (0.5 ng/ml, PBL), custom IFN-λ3 (20 ng/ml) and IFN-λ4 (50 ng/ml) (19), or were induced for 12 or 24 hours with dox (0.5 µg/ml), with untreated cells used as a negative control. Induction of ISRE-Luc reporter was evaluated by measuring Firefly and Renilla luciferase activity with dual-luciferase reporter assays on Glomax-Multi Detection System (Promega). The results were presented as relative light units (RLU), corresponding to Firefly normalized by Renilla luciferase activity.

### Interferon treatment

Cells were seeded in 12-well plates and treated for 8 hours in biological quadruplicates with IFNα (0.5 ng/ml, R&D Systems), IFNβ (0.5 ng/ml, GenScript), IFNγ (1 ng/ml, R&D Systems), IFN-λ1 (5 ng/ml, R&D Systems), IFN-λ2 (60 ng/ml, R&D Systems), and custom IFN-λ3 (20 ng/ml) and IFN-λ4 (50 ng/ml) (19), with no treatment used as a negative control.

### siRNA

For small interfering RNA (siRNA) knockdown, IFN-λ4-GFP HepG2 cells were induced with dox (0.1 µg/ml) and 24 hours later, transfected with a pool of siRNAs targeting *DDIT3* (Santa Cruz, sc-35437), *IRF1* (Silencer Select, ThermoFisher) or scrambled siRNA (Silencer Negative Control No. 1, ThermoFisher), using Lipofectamine RNAiMAX transfection reagent (ThermoFisher). Cells were harvested 72 hours after dox-induction and analyzed for DDIT3 (#5554, Cell Signaling Technology) and IRF1 (PA5-50512, ThermoFisher) protein expression by Western blotting.

### Western blotting

Cells were lysed in RIPA buffer (Sigma) supplemented with protease inhibitor cocktail (Promega) and PhosSTOP (Roche). Cell lysates were resolved on Blot 4-12% Bis-Tris gel (ThermoFisher) and transferred to nitrocellulose membranes with iBlot 2 Dry Blotting System (ThermoFisher). The membranes were probed with primary antibodies against IFNLR1 (#NBP1-84381, Novus Biologicals), STAT1 (#9172, Cell Signaling Technology), phospho-STAT1 (Tyr701, #58D6, Cell Signaling Technology), IFN-λ4 (ab196984; Abcam), GAPDH (ab37168, Abcam) and HRP-linked secondary antibody, goat anti-rabbit IgG (#7074; Cell Signaling Technology) or goat anti-mouse IgG (San Cruz, sc-2031). Signal detection was done with HyGLO™ quick spray chemiluminescent HRP antibody detection reagent (Denville Scientific) and Chemidoc Touch Imaging System (BioRad) and quantified by ImageJ software.

### Confocal microscopy

HepG2 cells were transfected for 24 hours with corresponding Halo-tagged constructs in 4-well chambered slides for fixed cells or in 4-well coverslip slides for live cells (2 × 10^5^ cells/well, LabTek). For fixed cells, a BacMam system (ThermoFisher) was used to deliver baculoviruses expressing GFP-tagged proteins targeted to specific organelles (N-acetylgalactosaminyltransferase for Golgi, Rab5a for early endosomes, Rab7a for late endosomes, and LAMP1 for lysosomes) 6 hours prior to transfection. Cells were incubated with cell-permeant TMR red Halo-tag ligand (1:2,000 for 15 min, Promega) 24 hours post-transfection, fixed with 4% paraformaldehyde, mounted with Prolong Gold antifade mounting media with DAPI (ThermoFisher) and coverslip.

For live cells, media in chambered slides was replaced with live-cell imaging solution (Life Technologies), supplemented with 20 mM glucose, 24 hours post-transfection. Cell-permeant TMR red Halo-tag ligand was added to cells (1:2,000 for 15 min). Imaging was done on an LSM700 confocal laser scanning microscope (Carl Zeiss) using an inverted oil lens at 40x magnification; live imaging was done using a temperature and CO2-controlled chamber, with 6 sec scans every minute for 12 hours.

### Tunicamycin treatment and Sendai Virus (SeV) infection

For SeV infection, PHH and HSC cells were infected with SeV (7.5 × 10^5^ chicken embryo ID50 [CEID50] per milliliter, Charles River Laboratories) and collected at the indicated time points. All experiments were done in triplicates.

### RNA extraction and quantitative reverse transcriptase-polymerase chain reaction (qRT-PCR) analysis

Total RNA was extracted using an RNeasy Mini Kit with on-column DNase digestion (Qiagen). RNA quantity and quality were evaluated by NanoDrop 8000 (Thermo Scientific) and Bioanalyzer 2100 (Agilent Technologies). RNA integrity numbers (RIN) of all RNA samples were > 9.5. cDNA was synthesized from the total RNA with the RT^2^ First Strand Kit (Qiagen), with an additional DNase I treatment step. qRT-PCR assays were performed in technical quadruplicates in 384-well plates on QuantStudio 7 instrument (Life Technologies), with RT^2^ SYBR Green (Qiagen) or TaqMan (Thermo Fisher) expression assays (**Table S8**). The expression of target genes was normalized by geometric means of endogenous controls (*GAPDH* and *ACTB*), presented as ΔCt values. The 2^-ΔΔCT^ method, in which ΔΔCt = ΔCt (experiment) – ΔCt (control), and fold = 2^-ΔΔCt^ was used to calculate the relative abundance of target mRNA expression and fold change. Data are shown as mean ± SEM based on biological replicates.

### Cell proliferation assays

For proliferation assays, HepG2 cells were induced with 0.5µg/ml dox and treated with 10 μM 5-bromo-2′-deoxyuridine (BrdU) for 3 h. Dead cells were detected with near-IR Live/Dead kit (ThermoFisher) and stained using a PE BrdU flow kit (BD Biosciences). For coculture experiments and proliferation experiments, HepG2 cells were labeled with Far Red dye (ThermoFisher). Cells were analyzed with multiparametric flow cytometry on a FACS Aria III (BD Biosciences) and FlowJo.v10 software (BD Biosciences).

### Cell viability, apoptosis and colony formation assays

Cells seeded in 96-well plates (8000 cells/well) were mock-or dox-induced (0.5 µg/ml) in quadruplicates for 24, 48 and 72 hours, then subjected to cell viability assays using CellTiter-Glo assays (Promega) and cell apoptosis assays using ApoTox-Glo Triplex assay (Promega). Luminescence was measured on the GloMax-multi detection system (Promega). For the colony formation assays, cells (2,000 cells/well) were mock- or dox-induced (0.5 µg/ml) in triplicates in 6-well plates for two weeks. The colonies were stained with 0.5% crystal violet, photographed and counted using ImageJ software. Cell cycle was evaluated using PI staining and analyzed on FlowJo.10 software (BD Biosciences). Seeded cells were fixed with 70% ethanol on ice for 2 hours, washed twice with PBS and treated with RNAse A (100µg/ml, Qiagen). Cells were then stained with PI (20µg/ml) for 10 minutes and were immediately analyzed on an Accuri C6 (BD Biosciences).

### RNA sequencing (RNA-seq)

Total RNA was extracted from cells using an RNeasy Mini Kit with an on-column DNase digestion (Qiagen) and quantitated using Qubit 4 Fluorometer (ThermoFisher). Ribosomal RNA was depleted with Ribo-Zero treatment, then 60 ng was used for library preparation using KAPA Stranded RNA-Seq Library Preparation Kit (Illumina, KR0934 – v 1.13). The samples were barcoded and pooled in sets of eight per run to generate paired 150-bp reads with NextSeq 550 Sequencing System (Illumina), generating 21.2 - 118.8 million reads per sample. The quality control analysis of RNA-seq data was carried out with FastQC pipeline. Paired RNA-seq reads were aligned to the ENSEMBL human reference genome GRCh37.75 (hg19) with STAR v 2.5.3a (49) using default parameters, and BAM files were generated with SAMtools v 1.5 (50). Data normalization and analysis of differential gene expression were done with DESeq2 R package v 1.22.2 (51, 52). Thresholds of fold change ≥ 1.5 (log2 fold change = 0.58) and false-discovery rate (FDR) adjusted P-value < 0.05 were used to identify differentially expressed genes (DEG) between experimental conditions. Identified DEGs were analyzed with Ingenuity Pathway Analysis (IPA) software (Qiagen). The generated RNA-seq dataset has been deposited into GEO NCBI (in progress).

### Statistical and bioinformatic analyses

#### Analysis of experimental data

Experimental data were analyzed using a two-sided, unpaired Student’s t-test, and P-values <0.05 were considered statistically significant. Statistical significance of the RNA-seq differential expression data was determined using Wald negative binomial test with Benjamini–Hochberg adjustment for multiple testing (52). Unless otherwise specified, data plotting and statistical analyses were performed with either Prism 7 (GraphPad), SPSS v.25 (IBM), or R packages. Means are presented with standard errors (SEM) based on biological replicates.

Inferring and analyzing TCGA-LIHC regulons enriched with expression signatures generated in HepG2 cells RNA-seq data (GRCh38/hg38) for the TCGA-HCC cohort (TCGA-LIHC, n=371) were downloaded from Genomic Data Commons using the R package TCGAbiolinks (53) and used to generate a transcriptional regulatory network containing regulons, each consisting of a transcription factor (TF) regulator and its inferred downstream targets. For regulators, we used the 1518 (94%) of 1612 TFs from (24) that were available in the TCGA-LIHC RNA-seq set.

We inferred the TCGA-BLCA regulons using the R package RTN version 2.10.1(23). Briefly, RNA-seq expression matrices for a set of samples were used to estimate the associations between a TF regulator and all of its potential targets. We used two metrics to identify potential regulator-target associations: Mutual Information (MI) and Spearman’s correlation. MI-based inference indicates whether a given regulator’s expression is informative of the expression of a given target gene, while a Spearman’s correlation indicates the positive or negative direction of an inferred association. Associations with MI below a minimum threshold were eliminated by permutation analysis (BH-adjusted p-value < 1×10^−7^), and unstable interactions were removed by bootstrapping, to create a regulatory network. Regulons were additionally processed by the ARACNe algorithm, which uses the data processing inequality (DPI) theorem to enrich the regulons with direct TF-target interactions (54).

The Master Regulator Analysis (MRA) assesses the overlap between a given regulon and a list of differentially expressed genes representing a given gene signature (55). The MRA was performed in R version 3.6.0 (R-Core-Team, 2012) using RTN (23).

We estimated regulon activity by a two-tailed gene set enrichment analysis (GSEA-2T), which is described elsewhere (23). Briefly, this approach assesses the distribution of regulon’s positive (A) and negative (B) targets over a list of genes ranked by a particular differential expression (DE) signature, which is called a ‘phenotype’. The distributions of A and B are then tested by GSEA statistics in the ranked phenotype, producing two independent enrichment scores (ES), and then a differential enrichment score (dES), which is obtained by subtracting the enrichment score for positive targets (ESA) from that obtained for negative targets (ESB). Large positive and negative dES indicate induced (activated) or repressed regulons. The statistical significance (nominal *P* value) of the *dES is estimated using an empirical phenotype-based permutation test procedure* (29). The GSEA-2T was performed in R (R-Core-Team, 2012) using the RTN package (23).

## ACKNOWLEDGMENTS

This study was supported by the Intramural Research Program of the Division of Cancer Epidemiology and Genetics, US National Cancer Institute, research grants from the Ministry of Science and Technology, Taipei, Taiwan (105-2628-B-010-003-MY4 and 107-2314-B-010-004-MY2). We thank the Cancer Genomics Research Laboratory (DCEG/NCI) for help with RNA-seq and cell lines authentication. We acknowledge all of the collaborators for REVEAL-II cohort: Hepatobiliary Division, Department of Internal Medicine, Kaohsiung Medical University, Kaohsiung, Taiwan: Chung-Feng Huang, Chia-Yen Dai, Jee-Fu Huang, Ming-Lun Yeh, Ching-I Huang, Ming-Lung Yu, Wan-Long Chuang; Division of Hepatogastroenterology, Department of Internal Medicine, China Medical University Hospital, Taichung, Taiwan: Hsueh-Chou Lai, Wen-Pang Su, Jung-Ta Kao, Sheng-Hung Chen, Po-Heng Chuang, Cheng-Yuan Peng; Department of Gastroenterology and Hepatology, Linkou Medical Center, Chang Gung Memorial Hospital, Kweishan, Taoyuan, Taiwan:Chun-Yen Lin, Wen-Juei Jeng, I-Shyan Sheen; Department of Internal Medicine, National Taiwan University Hospital and National Taiwan University College of Medicine, Taipei, Taiwan: Chen-Hua Liu, Chun-Jen Liu, Hung-Chih Yang, Chieh-Chang Chen, Shih-Jer Hsu, Jia-Horng Kao; Division of Hepato-Gastroenterology, Department of Internal Medicine, Kaohsiung Chang Gung Memorial Hospital and Chang Gung University College of Medicine, Kaohsiung, Taiwan: Jing-Houng Wang, Kwong-Ming Kee, Sheng-Nan Lu; Academia Sinica, Taipei, Taiwan: Hwai-I Yang, Chien-Jen Chen; Global Health Economics and Outcomes Research, Bristol Myers-Squibb, Princeton, NJ, USA:Yong Yuan. The presented results are in part based upon data generated by the TCGA Research Network.

