## Supplementary Material for "IFN-λ4 may contribute to HCV persistence by increasing ER stress and enhancing IRF1 signaling"

### SUPPLEMENTARY MATERIALS

#### Supplementary Tables

**Table S1.** Baseline characteristics of HCV-infected patients in the REVEAL II cohort, Taiwan

**Table S2.** Association of the *IFNL4* genotype with reduced SVR in HCV-infected patients in the REVEAL II cohort, Taiwan

**Table S3.** Progression to HCC in HCV-infected patients in the REVEAL II cohort, Taiwan in relation to *IFNL4* genotype and SVR after treatment/retreatment with peg-IFNα/RBV

**Table S4.** Progression to HCC in HCV-infected patients in the REVEAL II cohort, Taiwan in relation to *IFNL4* genotype and cirrhosis

**Table S5.** *IFNL4* genotype is not associated with an increased risk of HCC in HBV-infected patients in China

**Table S6.** List of DEGs significantly induced by IFN-λ4-GFP, separate Excel file

**Table S7.** List of regulons derived from DEGs significantly induced by IFN-λ4-GFP, separate Excel file

**Table S8.** Primers and expression assays used in the study

#### Supplementary Figures

**Figure S1.** Generation of the IFN-λ4-GFP-IFNLR1$^{KO}$ HepG2 cell line using CRISPR/Cas9 genome editing

**Figure S2.** IFNLR1 knock-out in IFN-λ4-GFP-IFNLR1$^{KO}$ HepG2 cells eliminates the induction of the JAK/STAT signaling by type III IFNs without affecting signaling of other IFNs

**Figure S3.** Outline of RNA-seq data analysis in HepG2 cells

**Figure S4.** Comparison of *IFNL3* and *IFNL4* mRNA expression in HepG2 cell lines

**Figure S5.** Top 50 IFNLR1-dependent ISGs induced by expression of IFN-λ3-GFP and IFN-λ4-GFP in HepG2 cells

**Figure S6.** IFNLR1-dependent pathways affected by IFN-λ4-GFP expression in HepG2 cells

**Figure S7.** IFN-λ4 expression significantly inhibits proliferation in HepG2 cells

**Figure S8.** *IRF1* is upregulated by IFN-λ4-GFP expression in HepG2 cells

**Figure S9.** Pathways for IFNLR1-dependent DEGs significantly affected by IFN-λ4-GFP expression in HepG2 cells

**Figure S10.** IFN-λ4 expression induces signaling pathways related to unfolded protein response (UPR).

**Figure S11.** *IFNL4* genotype is associated with increased ER stress in primary human hepatocytes (PHH) infected with Sendai virus (SeV)

### Supplementary Videos

**Video S1**. Trafficking of IFN-λ4 in late endosomes in HepG2 cells

**Video S2**. Apoptosis of HepG2 cells expressing IFN-λ4

### SUPPLEMENTARY TABLES

**Table S1. Baseline characteristics of HCV-infected patients in the REVEAL II cohort, Taiwan**

| Characteristics | Total N=2931 | Liver cirrhosis at baseline | | P-value[#] |
|---|---|---|---|---|
| | | Yes N=508 (SD or %) | No N=2423 (SD or %) | |
| Age (years) | 53.46 (9.81) | 56.03 (8.68) | 52.92 (9.95) | <0.0001 |
| Follow-up (years) | 6.19 (4.33) | 6.55 (4.06) | 6.12 (4.39) | 0.030 |
| ALT, U/L | 127.7 (101.1) | 132.9 (94.54) | 126.6 (102.3) | 0.188 |
| Platelet count, $10^3$/µL | 150.1 (78.10) | 116.8 (109.6) | 156.8 (153.5) | <0.0001 |
| Gender | | | | 0.0004 |
| Male | 1455 (49.64) | 216 (42.52) | 1239 (51.13) | |
| Female | 1476 (50.36) | 292 (57.48) | 1184 (48.87) | |
| *IFNL4*-rs368234815 | | | | 0.044 |
| TT/TT | 2618 (89.32) | 468 (92.13) | 2150 (88.73) | |
| TT/dG | 303 (10.34) | 40 (7.87) | 263 (10.85) | |
| dG/dG | 10 (0.34) | 0 (0.00) | 10 (0.41) | |
| SVR at initial treatment | | | | <0.0001 |
| No | 568 (19.38) | 144 (29.21) | 424 (18.07) | |
| Yes | 2271 (77.48) | 349 (70.79) | 1922 (81.93) | |
| Unknown | 92 (3.14) | | | |
| SVR at any treatment* | | | | <0.0001 |
| Never | 454 (15.49) | 112 (22.72) | 342 (14.58) | |
| Ever | 2385 (81.37) | 381 (77.28) | 2004 (85.42) | |
| Unknown | 92 (3.14) | | | |
| HCV genotype[$] | | | | 0.871 |
| Genotype 1 | 1625 (55.44) | 280 (55.12) | 1345 (55.51) | |
| Other | 1306 (44.56) | 228 (44.88) | 1078 (44.49) | |

[#]T-test for continuous variables and Chi-squared test for categorical variables; *HCV genotype 1 vs. other genotypes (2, 3, 4, and 6); [$] SVR – sustained virologic response either at initial treatment or retreatment.

**Table S2. Association of the *IFNL4* genotype with reduced SVR in HCV-infected patients in the REVEAL II cohort, Taiwan**

| Genotype of *IFNL4*-rs368234815 | SVR, N=2839 N (%) | | OR (95% CI), P-value Covariates in multivariable models | | | | | |
|---|---|---|---|---|---|---|---|---|
| | Yes | No | crude | age, sex | age, sex, ALT | age, sex, baseline cirrhosis, ALT | HCV genotype 1 age, sex, baseline cirrhosis, ALT | other HCV genotypes* age, sex, baseline cirrhosis, ALT |
| **Initial Treatment with peg-IFNα/RBV** | | | | | | | | |
| TT/TT | 2091 (82.16) | 454 (17.84) | Ref | Ref | Ref | Ref | Ref | Ref |
| TT/dG and dG/dG | 180 (61.22) | 114 (38.78) | 0.343 (0.265-0.443) 2.24E-16 | 0.343 (0.265-0.443) 2.47E-16 | 0.354 (0.272-0.459) 5.62E-15 | 0.338 (0.260-0.440) 7.33E-16 | 0.310 (0.225-0.427) 7.43E-13 | 0.462 (0.264-0.808) 0.007 |
| **Initial Treatment and Retreatment with peg-IFNα/RBV** | | | | | | | | |
| TT/TT | 2193 (86.17) | 352 (13.83) | Ref | Ref | Ref | Ref | Ref | Ref |
| TT/dG and dG/dG | 192 (65.31) | 102 (34.69) | 0.302 (0.232-0.394) 9.11E-19 | 0.302 (0.231-0.395) 1.63E-18 | 0.318 (0.242-0.418) 1.68E-16 | 0.308 (0.234-0.405) 3.60E-17 | 0.265 (0.191-0.368) 1.66E-15 | 0.560 (0.296-1.060) 0.075 |

*Other HCV genotypes include genotypes 2, 3, 4, and 6; SVR – sustained virologic response; ALT - alanine aminotransferase.

**Table S3. Progression to HCC in HCV-infected patients in the REVEAL II cohort, Taiwan in relation to *IFNL4* genotype and SVR after treatment/retreatment with peg-IFNα/RBV**

| | | | | HR (95% CI), P-value<br>Covariates in multivariable models | | | | |
|---|---|---|---|---|---|---|---|---|
| Genotype of *IFNL4*-rs368234815 | HCC, N (%) | Total N, person-year of follow-up | Incidence (1/1000) | Crude | age, sex | age, sex, baseline cirrhosis | age, sex, baseline cirrhosis, ALT | age, sex, baseline cirrhosis, ALT, HCV genotype# |
| **HCV patients with SVR information, n=2839*** | | | | | | | | |
| TT/TT | 109 (84.50) | 2545 (15542.24) | 7.01 | Ref | Ref | Ref | Ref | Ref |
| TT/dG and dG/dG | 20 (15.50) | 294 (1725.00) | 11.59 | 1.71 (1.06-2.75) **P=0.028** | 1.58 (0.98-2.56) P=0.060 | 1.69 (1.05-2.73) **P=0.033** | 1.74 (1.07-2.83) **P=0.024** | 1.71 (1.05-2.77) **P=0.030** |
| **Ever SVR - at initial treatment or retreatment, N=2385** | | | | | | | | |
| TT/TT | 78 | 2193 (13329.75) | 5.85 | Ref | Ref | Ref | Ref | Ref |
| TT/dG or dG/dG | 7 | 192 (1171.57) | 5.97 | 1.05 (0.49-2.28) P=0.90 | 0.98 (0.45-2.13) P=0.96 | 1.10 (0.51-2.40) P=0.81 | 1.14 (0.52-2.49) P=0.74 | 1.14 (0.52-2.49) P=0.74 |
| **Never SVR - at initial treatment or retreatment, N=454** | | | | | | | | |
| TT/TT | 31 | 352 (2212.49) | 14.01 | Ref | Ref | Ref | Ref | Ref |
| TT/dG or dG/dG | 13 | 102 (553.42) | 23.49 | 1.77 (0.92-3.40) P=0.088 | 1.66 (0.86-3.18) P=0.13 | 1.82 (0.94-3.53) P=0.075 | 1.87 (0.96-3.64) P=0.065 | 1.81 (0.93-3.53) P=0.081 |

*SVR status was unknown for 92 HCV patients; SVR – sustained virologic response; ALT - alanine aminotransferase; # HCV genotype 1 vs. other genotypes (2, 3, 4, and 6).

**Table S4. Progression to HCC in HCV-infected patients in the REVEAL II cohort, Taiwan in relation to *IFNL4* genotype and cirrhosis**

| | | | | HR (95% CI), P-value<br>Covariates in multivariable models | | | | |
|---|---|---|---|---|---|---|---|---|
| Genotype of *IFNL4*-rs368234815 | HCC, N (%) | Total N, person-year of follow-up | Incidence (1/1000) | Crude | age, sex | age, sex, SVR at any treatment[#] | age, sex, ALT SVR at any treatment | age, sex, ALT, HCV genotype[#], SVR at any treatment |
| **All HCV patients with information on baseline cirrhosis status, n=2931 (100%)** | | | | | | | | |
| TT/TT | 111 (82.84) | 2618 (16279.35) | 6.82 | Ref | Ref | Ref | Ref | Ref |
| TT/dG or dG/dG | 23 (17.16) | 313 (1816.60) | 12.66 | 1.91 (1.22-2.99) P=0.0049 | 1.79 (1.14-2.82) P=0.011 | 1.38 (0.85-2.25) P=0.195 | 1.45 (0.89-2.36) P=0.138 | 1.45 (0.89-2.36) P=0.141 |
| **HCV patients with baseline cirrhosis, N=508 (17.3%)** | | | | | | | | |
| TT/TT | 51 | 468 (3127.33) | 16.31 | Ref | Ref | Ref | Ref | Ref |
| TT/dG or dG/dG | 10 | 40 (197.71) | 50.58 | 3.30 (1.66-6.56) P=0.0007 | 2.32 (1.15-4.68) P=0.019 | 1.83 (0.88-3.79) P=0.103 | 2.00 (0.96-4.16) P=0.066 | 2.00 (0.96-4.16) P=0.066 |
| **HCV patients without baseline cirrhosis, N=2423 (82.7%)** | | | | | | | | |
| TT/TT | 60 | 2150 (13152.02) | 4.56 | Ref | Ref | Ref | Ref | Ref |
| TT/dG or dG/dG | 13 | 273 (1621.89) | 8.02 | 1.80 (0.99-3.27) P=0.055 | 1.80 (0.99-3.29) P=0.055 | 1.22 (0.62-2.41) P=0.57 | 1.25 (0.63-2.48) P=0.52 | 1.25 (0.63-2.47) P=0.53 |

[#] SVR – sustained virologic response either at initial treatment or retreatment with peg-IFNα/RBV; *HCV genotype 1 vs. other genotypes (2, 3, 4, and 6).

**Table S5.** ***IFNL4*** **genotype is not associated with an increased risk of HCC in HBV-infected patients in China**

| Genotypes of *IFNL4*-rs368234815 | HBV, HCC patients n=1291 N (%) | HBV persistent n=1332 N (%) | HBV spontaneous clearance n=1320 N (%) | OR (95% CI)[a] | *P* [a] | OR (95% CI)[b] | *P* [b] |
|---|---|---|---|---|---|---|---|
| TT/TT | 1128 (87.4) | 1158 (86.9) | 1163 (88.1) | Ref | | Ref | |
| dG/TT | 156 (12.1) | 167 (12.5) | 152 (11.5) | 0.95 (0.75-1.21) | 0.694 | 1.10 (0.87-1.39) | 0.424 |
| dG/dG | 7 (0.5) | 7 (0.5) | 5 (0.4) | 0.91 (0.31-2.63) | 0.857 | 1.42 (0.45-4.50) | 0.548 |
| Dominant TT/TT vs. dG/TT&dG/dG | 12.6 | 12.7 | 11.9 | 0.95 (0.76-1.20) | 0.676 | 1.11 (0.88-1.40) | 0.372 |
| Additive, per allele | | | | | | | |
| TT | 93.4 | 93.2 | 93.9 | Ref | | Ref | |
| dG | 6.6 | 6.8 | 6.1 | 0.95 (0.77-1.19) | 0.669 | 1.11 (0.89-1.38) | 0.338 |

Logistic regression analyses adjusted for age, gender, smoking and drinking status.
[a]HCC patients vs. HBV persistent carriers; [b]HBV persistent carriers vs. HBV natural clearance subjects.

### Table S8. Primers and expression assays used in the study

| | Primer or gene | Primer sequence (5'-3') or gene assays |
|---|---|---|
| **IFNLR1** | Exon3-F | CAAAGCTAGGGAGGGACAGGG |
| **Primers** | Exon3-R | CTGACAGGCTACCAGACATG |
| | Sequencing | GTCATCAACCCGCTCCAAGG |
| **Taqman gene expression assays** (ThermoFisher Scientific) | | |
| | *GAPDH* | Hs02786624_g1 |
| | *ACTB* | 4326315E |
| | *MX1* | Hs00895608_m1 |
| | *ISG15* | ISG15 Hs01921425_s1 |
| | *OAS1* | Hs00973637_m1 |
| | *CXCL10* | Hs00171042_m1 |
| | *PD-L1* | Hs01125301_m1 |
| | *PERK* | Hs00984005_m1 |
| | *ATF4* | Hs00909569_g1 |
| | *ATF6* | Hs00232586_m1 |
| | *DDIT3* | Hs00358796_g1 |
| | *FGF21* | Hs00173927_m1 |
| | *VLDLR* | Hs01045914_g1 |
| | *VLDLRAS1* | Hs03309901_g1 |
| | *IRF1* | Hs00971965_m1 |
| | *COL1A1* | Hs00164004_m1 |
| **Custom Taqman gene expression assay** (ThermoFisher Scientific) | | |
| | *XBP1S* | |
| | Forward primer | GCTGAGTCCGCAGCAGGT |
| | Probe | CCAACAGGATATCAGACTCTGAATCT |
| | Reverse primer | CAGAACATCTCCCCATGGA |

### SUPPLEMENTARY FIGURES

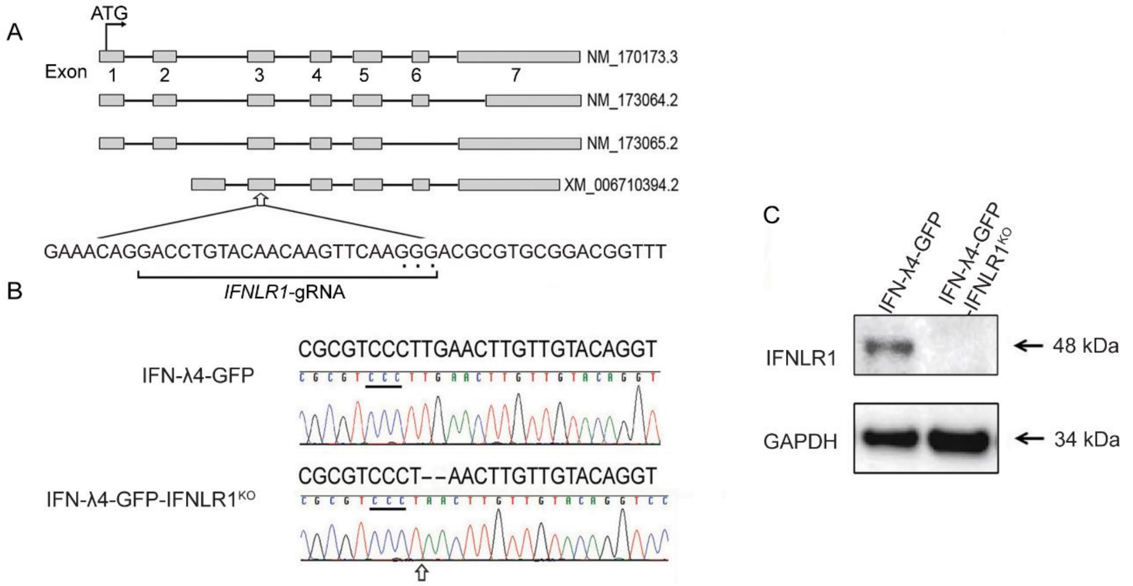


**Fig. S1. Generation of the IFN-λ4-GFP-IFNLR1$^{KO}$ HepG2 cell line using CRISPR/Cas9 genome editing.**

All type III IFNs signal via a heterodimeric receptor complex composed of the ubiquitously expressed interleukin 10 receptor 2 (IL10R2) that is shared by many cytokines, and the interferon lambda receptor 1 (IFNLR1), which is primarily expressed in epithelial and hepatic cells and serves only type III IFNs. Both receptors are expressed in HepG2, a hepatoma cell line, which responds to the signaling of type III IFNs. The HepG2-IFNLR1$^{KO}$ cell line was created by eliminating *IFNLR1* in HepG2 cells by CRISPR-Cas9 genome editing. **(A)** Genomic organization of the *IFNLR1* transcripts. Arrow indicates the region within the common exon 3 targeted by all six *IFNLR1*-gRNAs tested, with exon numbering based on the NM_170173.3 transcript. Target sites and protospacer adjacent motif (PAM) sequences for the most efficient *IFNLR1*-gRNA are marked by lines and dots, respectively. **(B)** Sequence alignment of the target region of *IFNLR1* exon 3 in the IFN-λ4-GFP and IFN-λ4-GFP-IFNLR1$^{KO}$ HepG2 cell lines shows a 2 bp deletion that is expected to eliminate the IFNLR1 protein. **(C)** Complete elimination of the IFNLR1 protein is validated by Western blotting of cell lysates, with GAPDH used as a loading control. Top ten predicted off-target sites were tested by sequencing of the genomic DNA of IFN-λ4-GFP-IFNLR1$^{KO}$ cells and no off-target mutations were detected (data not shown).

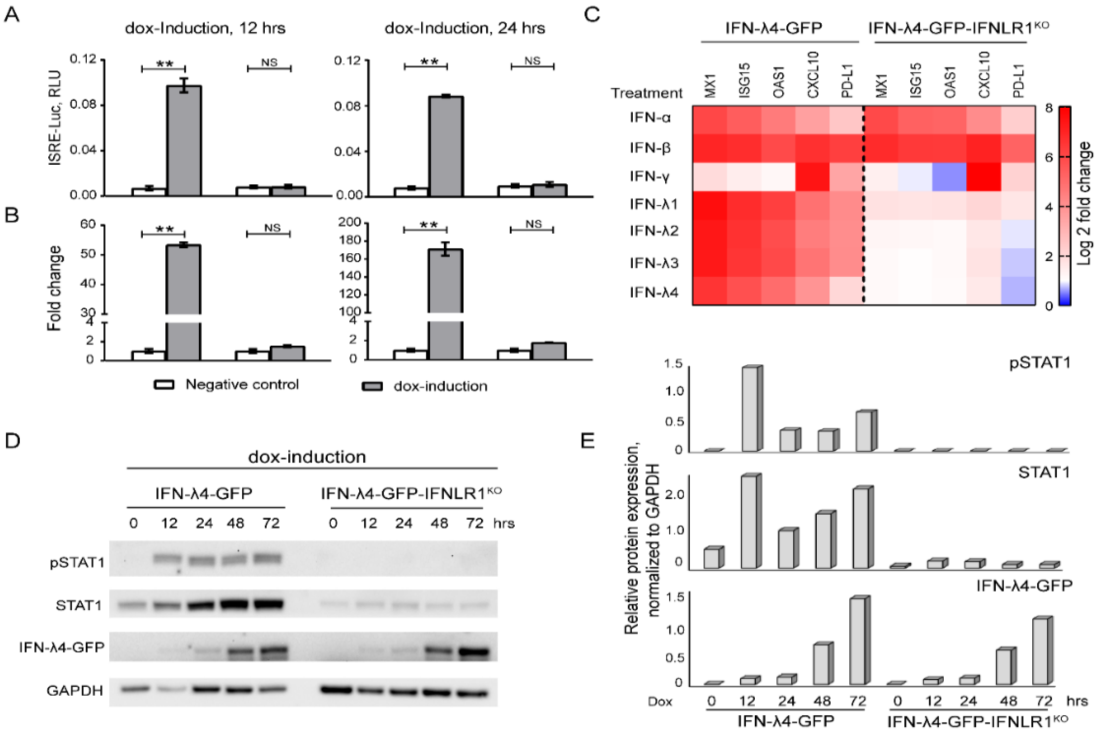


**Fig. S2. IFNLR1 knock-out in IFN-λ4-GFP-IFNLR1$^{KO}$ HepG2 cells eliminates the induction of the JAK/STAT signaling by type III IFNs without affecting signaling of other IFNs.**

**(A)** IFN-λ4-GFP and IFN-λ4-GFP-IFNLR1$^{KO}$ HepG2 cells were transiently transfected in 96-well plates with the Cignal ISRE-Luc reporter plasmid and dox-induced for 12 or 24 hrs; uninduced cells were used as negative controls. Activation of the JAK/STAT signaling was evaluated by the ability to induce the ISRE-Luc reporter, measured in RLU (relative luciferase units), all based on biological triplicates per condition. **(B)** qRT-PCR expression analysis of *ISG15* with *ACTB* used as an endogenous control; the results are normalized to the control group, based on biological duplicates. **(C)** IFN-λ4-GFP and IFN-λ4-GFP-IFNLR1$^{KO}$ cells were treated in 4 biological replicates with recombinant human proteins - IFNα (0.5 ng/ml), IFNβ (0.5 ng/ml), IFNγ (1 ng/ml), IFN-λ1 (5 ng/ml), IFN-λ2 (60 ng/ml), IFN-λ3 (20 ng/ml) and IFN-λ4 (50 ng/ml) for 8 hrs, with an untreated group used as control. Expression of *MX1*, *ISG15*, *OAS1*, *CXCL10* and *PD-L1* was analyzed by qRT-PCR in 4 technical replicates. Heatmap shows the expression of each gene analyzed in ddCt (log2 fold change) values between treated and control groups. **(D)** Western blotting of pSTAT1, STAT1 and IFN-λ4-GFP expression in whole-cell lysates of corresponding cells, with GAPDH expression used as a loading control. **(E)** Results of Image J quantitative analysis of Western blots of pSTAT1, STAT1 and IFN-λ4-GFP expression (from panel **D**), all normalized to GAPDH. As expected, type III IFN signaling in the IFN-λ4-GFP-IFNLR1$^{KO}$ was completely abrogated without affecting signaling of other IFNs that do not signal through IFNLR1. Error bars – SEM; ** - $P < 0.01$; NS – not significant based on two-sided Student's T-test.

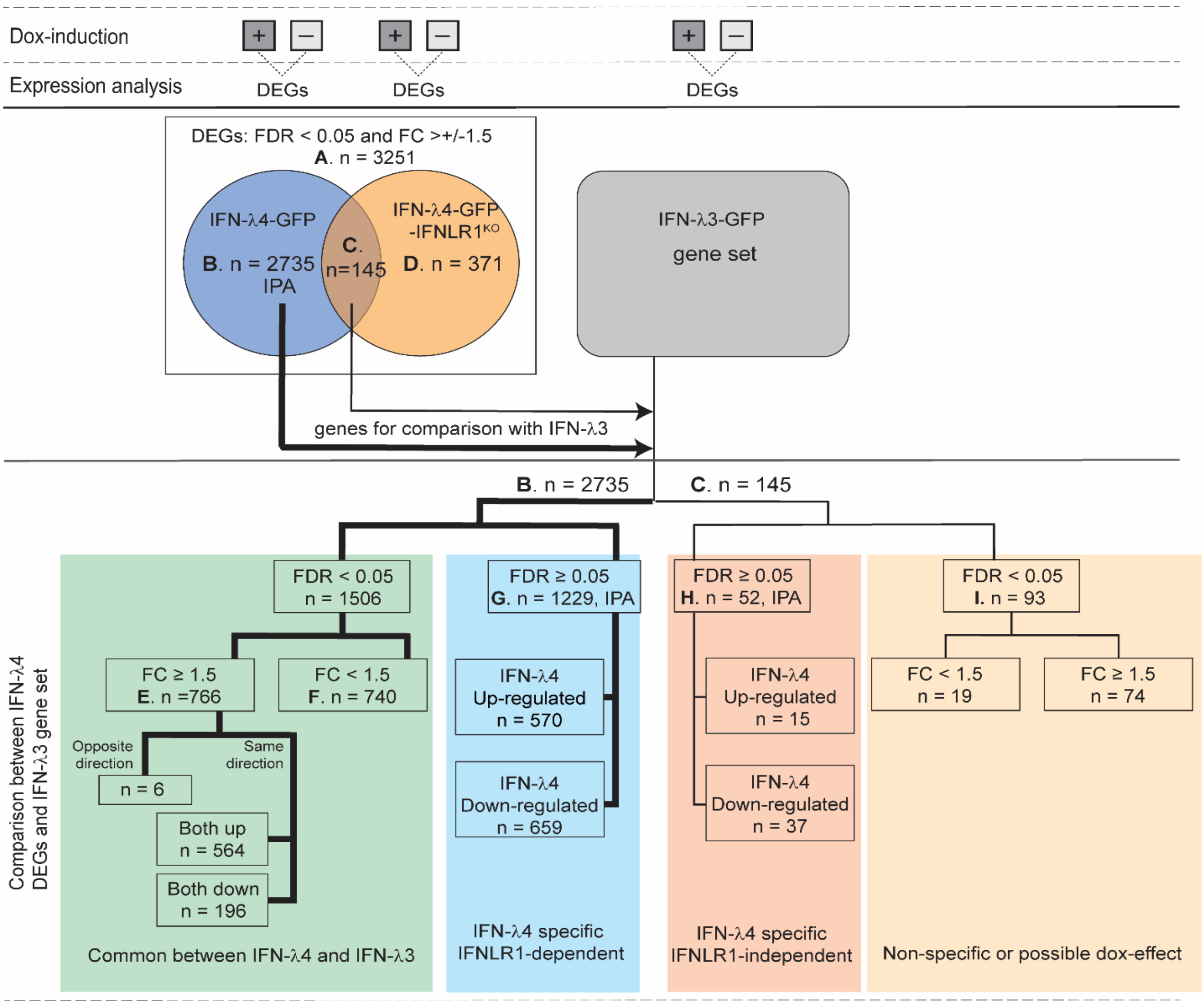


**Fig. S3. Outline of RNA-seq analysis in HepG2 cells**.

RNA-sequencing was performed using total RNA from IFN-λ4-GFP, IFN-λ4-GFP-IFNLR1$^{KO}$ and IFN-λ3-GFP HepG2 cells in biological triplicates. Differentially expressed genes (DEGs, FDR<0.05, +/- ≥1.5-fold change (FC)) were identified by comparing corresponding dox+ and dox- cells after 72 hrs of induction. **(A)** All DEGs (n=3,251) induced by IFN-λ4-GFP; **(B)** DEGs (n=2,735) induced by IFN-λ4-GFP and IFNLR1-dependent; **(C)** DEGs (n=145) induced by IFN-λ4-GFP and IFNLR1-independent; **(D)** DEGs (n=371) induced only in IFN-λ4-GFP-IFNLR1$^{KO}$. DEGs unique to IFN-λ4-GFP (thick line) and those common to both IFN-λ4-GFP and IFN-λ4-GFP-IFNLR1$^{KO}$ (thin line) were analyzed in the transcriptome of IFN-λ3-GFP cells; **(E)** DEGs (n=766) also significant in IFN-λ3-GFP (FDR<0.05, ≥1.5 FC); **(F)** DEGs (n=740) also significant (FDR<0.05) but with <1.5 FC **(G)** DEGs (n=1,229) considered IFN-λ4-GFP-specific and IFNLR1-dependent; **(H)** DEGs (n=52) considered IFN-λ4-GFP-specific and IFNLR1-independent; **(I)** DEGs (n=93) non-specifically induced in all 3 cell lines. IPA – sets of genes used for Ingenuity Pathway, additional details are provided in **Table S6.**

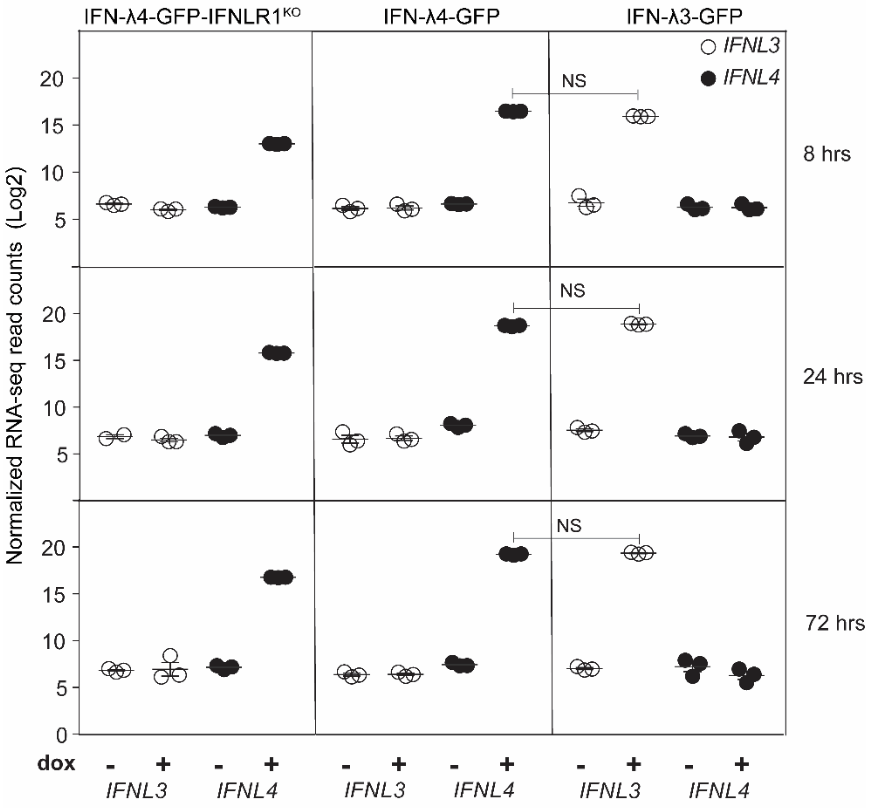


**Fig. S4. Comparison of *IFNL3* and *IFNL4* mRNA expression in HepG2 cell lines.**

Normalized counts of *IFNL3* and *IFNL4* RNA-seq reads (Log2) in corresponding cell lines with/without dox induction for indicated time points. *IFNL3* and *IFNL4* were induced specifically and to a similar magnitude. Shown - individual and mean values for biological triplicates. NS - not statistically significant based on two-sided Student's T-test.

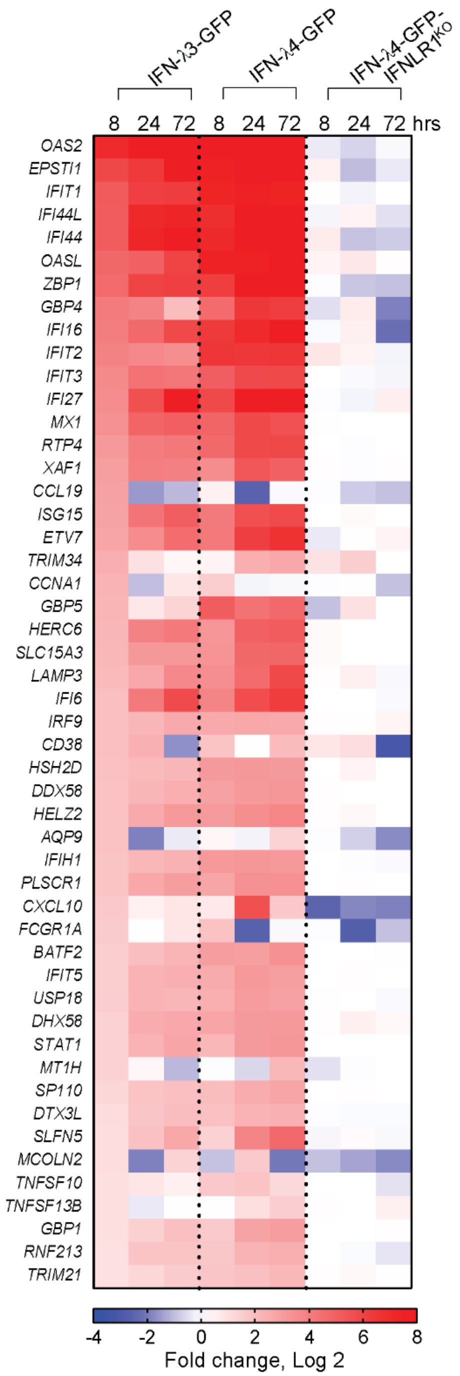


**Fig. S5. Top 50 IFNLR1-dependent ISGs induced by expression of IFN-λ3-GFP and IFN-λ4-GFP in HepG2 cells.**

Differentially expressed genes (DEGs) were identified by RNA-seq analysis in IFN-λ3-GFP, IFN-λ4-GFP or IFN-λ4-GFP-IFNLR1$^{KO}$ HepG2 cells, comparing corresponding dox+/dox-cells. Heatmap shows 50 most significantly induced IFNLR1-dependent ISGs presented as fold change (Log 2) at indicated time points; additional details are provided in **Table S6.**

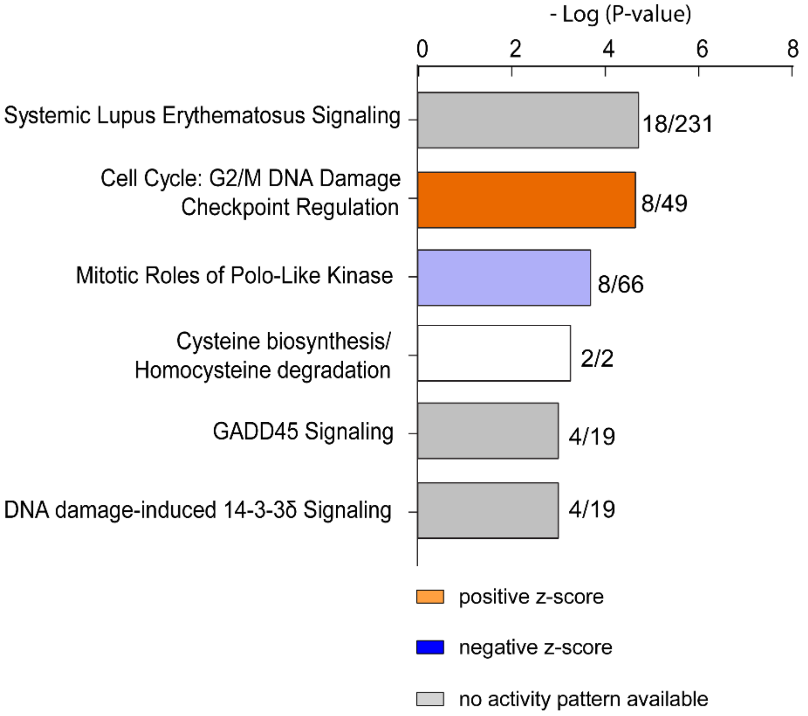


**Fig. S6. IFNLR1-dependent pathways affected by IFN-λ4-GFP expression in HepG2 cells.**

Ingenuity Pathway Analysis (IPA) of 1,229 DEGs (**Fig. S3G**) in IFN-λ4-GFP HepG2 cells after 72 hrs of dox induction (P-FDR < 0.05, fold change (Log2) > +/-1.5). P-values (-log) represent the significance of gene set enrichment in each pathway.

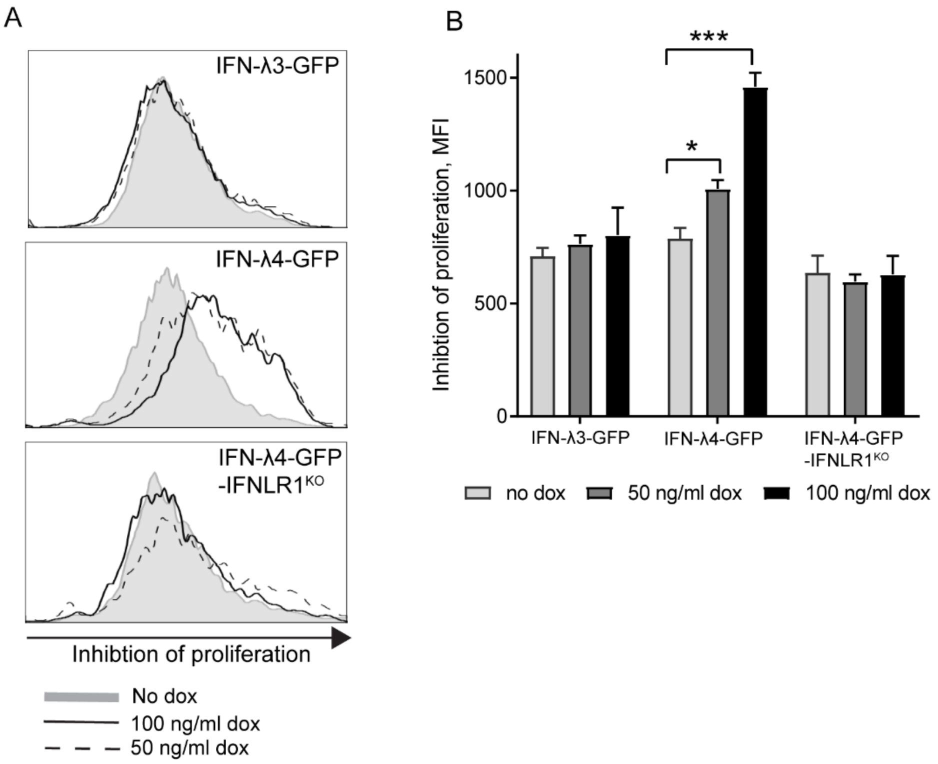


**Fig. S7. IFN-λ4 expression significantly inhibits proliferation in HepG2 cells.**

**(A)** IFN-λ3-GFP, IFN-λ4-GFP and IFN-λ4-GFP-IFNLR1$^{KO}$ HepG2 cells were labeled with the Far Red dye to monitor cell proliferation. The cells were dox-induced for 5 days at indicated concentrations and proliferation was assessed by flow cytometry, with results presented as histograms. **(B)** Higher Far Red mean fluorescence intensity values indicate proliferation inhibition. Shown – one representative of three independent experiments. Error bars – SEM, based on 3 biological replicates. P-values are for comparisons between corresponding dox+ and dox- cells using the two-sided Student's T-test. * $p<0.05$, *** $p<0.001$.

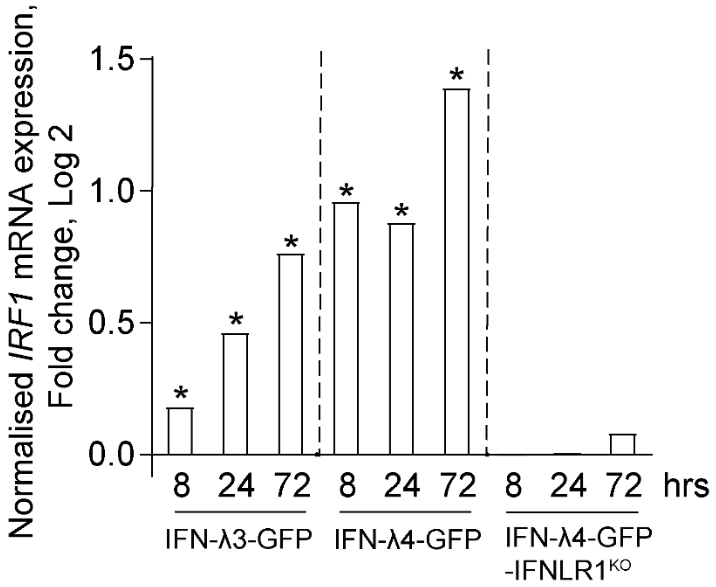


**Fig. S8.** ***IRF1*** **is upregulated by IFN-λ4-GFP expression in HepG2 cells.**

*IRF1* mRNA expression is presented as fold change (Log2) between corresponding dox+ and dox- HepG2 cells (based on RNA-seq data, **Table S6)**. *P-FDR < 0.05.

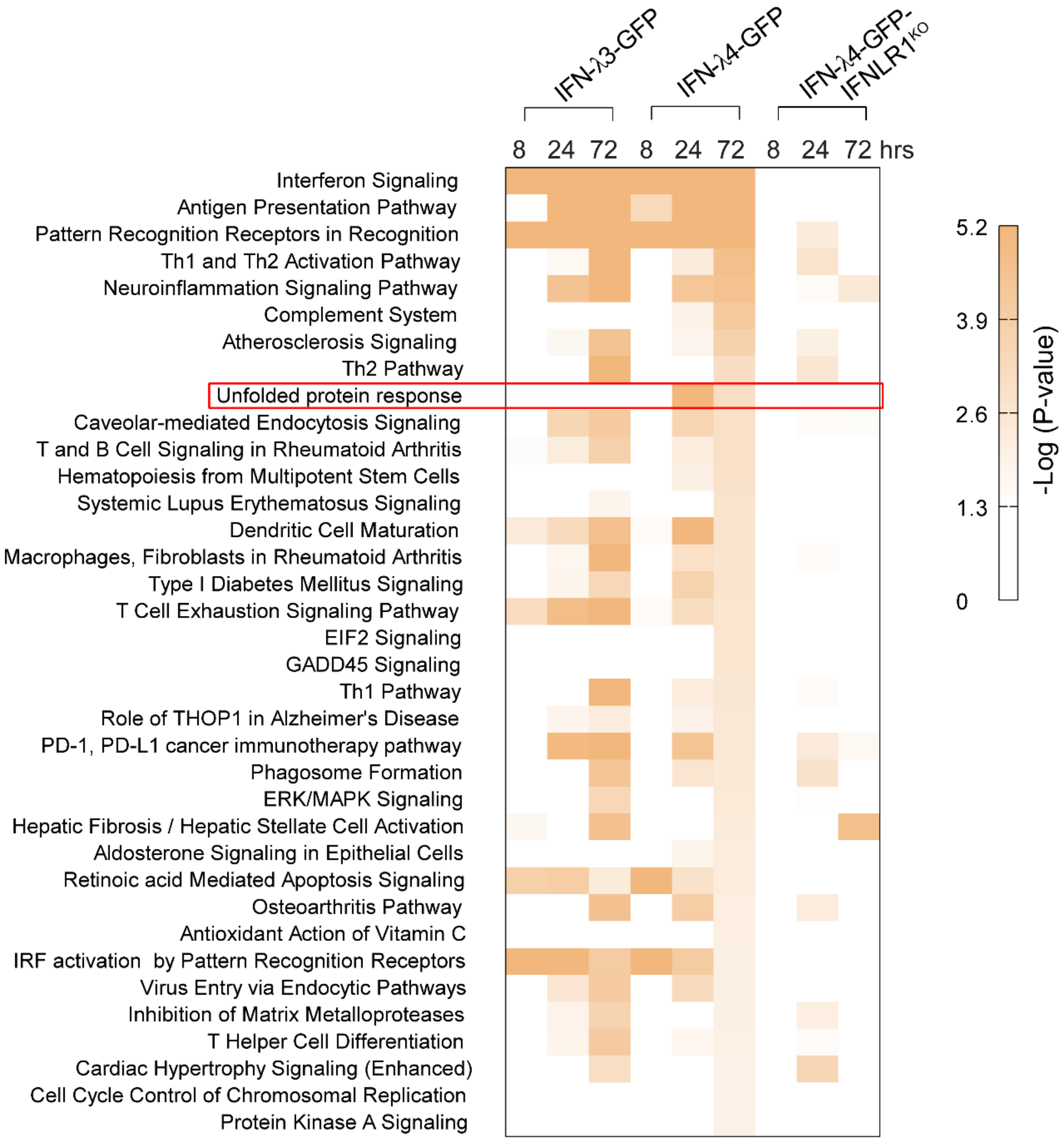


**Fig. S9. Pathways for IFNLR1-dependent DEGs significantly affected by IFN-λ4-GFP expression in HepG2 cells.**

Ingenuity Pathway Analysis (IPA) of differentially expressed genes (DEGs, P-FDR < 0.05, fold change > +/-1.5) identified by RNA-seq in IFN-λ3-GFP, IFN-λ4-GFP and IFN-λ4-GFP-IFNLR1$^{KO}$ HepG2 cells comparing corresponding dox+ and dox- cells after 72 hrs of induction. All IFNLR1-dependent DEGs of IFN-λ4-GFP (n=2735, **Fig. S3B**) were used as input. Enriched pathways identified in IFN-λ4-GFP cells (P < 0.01) were plotted in a heatmap for all 3 cell lines with intensity (brown) representing (-log) P-value of the significance level of gene set enrichment. Unfolded protein response (UPR) pathway is marked as uniquely induced by IFN-λ4-GFP and not IFN-λ3-GFP.

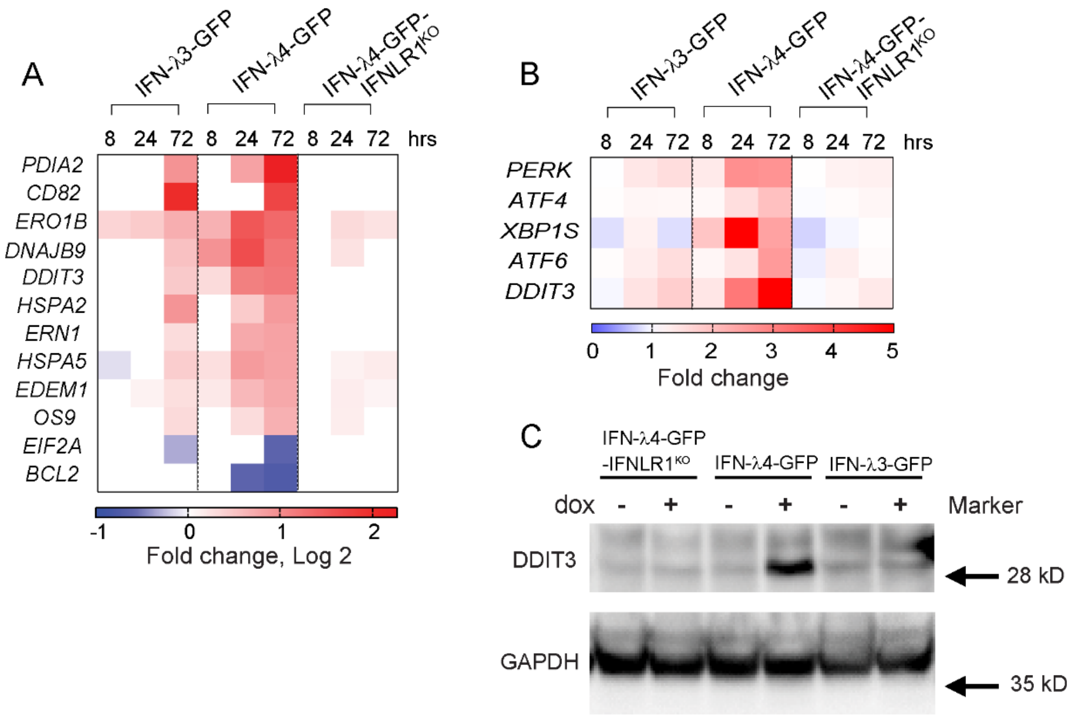


**Fig. S10. IFN-λ4 expression induces signaling pathways related to unfolded protein response (UPR).**

**(A)** RNA-seq data of select differentially expressed UPR-related genes in HepG2 cells after dox-induction for indicated time points. **(B)** Heatmap of induction of several UPR-related genes based on qRT-PCR analysis in an independent set of HepG2 samples, comparing dox+ vs dox- conditions. **(C)** Western blot showing DDIT3 protein expression comparing corresponding dox+ and dox- HepG2 cells induced for 72 hrs. GAPDH is used as a loading control.

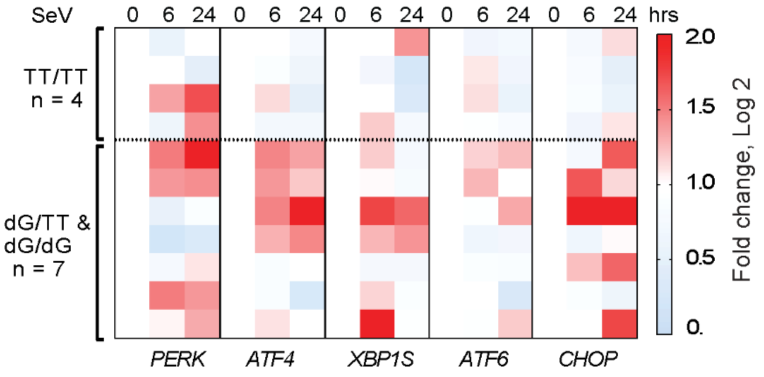


**Fig. S11. *IFNL4* genotype is associated with increased ER stress in primary human hepatocytes (PHH) infected with Sendai virus (SeV).**

Endoplasmic reticulum (ER) stress is evaluated by the expression of select unfolded protein response (UPR) genes in PHH infected with SeV for 24 hrs. Expression was measured with qRT-PCR assays and normalized to an endogenous control (*GAPDH*) and uninfected cells. Results are presented as a heatmap (fold change, Log2), according to *IFNL4* genotype groups.

### Supplementary Videos

**Video S1**. Trafficking of IFN-λ4 in late endosomes in HepG2 cells.

HepG2 cells were transiently transfected with expression constructs IFN-λ4-Halo and GFP-Rab7a (late endosome marker). Cells were stained with a cell-permeant TMR Halo-tag ligand (red) and live-imaged starting 24 hrs post-transfection. Images were collected every minute for a total of 12 hrs. Snapshots of this video are presented in **Fig. 3B**. Video length: 12 sec

**Video S2**. Apoptosis of HepG2 cells expressing IFN-λ4.

HepG2 cells were transiently transfected with the IFN-λ4-Halo expression construct and live-imaged as described in **Video S1**. Video shows significant membrane blebbing immediately before cell rupture. Snapshots of this video are presented in **Fig. 3D**. Video length: 80 sec
